# A thermodynamic chemical reaction network drove autocatalytic prebiotic peptides formation

**DOI:** 10.1101/461707

**Authors:** Peng Bao, Yu-Qin He, Guo-Xiang Li, Hui-En Zhang, Ke-Qing Xiao

## Abstract

The chemical reaction networks (CRNs), which led to the transition on early Earth from geochemistry to biochemistry remain unknown. We show that under mild hydrothermal circumstances, a thermodynamic chemical reaction network including sulfite/sulfate coupled with anaerobic ammonium oxidation (Sammox), might have driven prebiotic peptides synthesis. Peptides comprise 14 proteinogenic amino acids, endowed Sammox-driven CRNs with autocatalysis. The peptides exhibit both forward and reverse catalysis, with the opposite catalytic impact in sulfite- and sulfate-fueled Sammox-driven CRNs, respectively, at both a variable temperature range and a fixed temperature, resulting in seesaw-like catalytic properties. The ratio of sulfite to sulfate switches the catalytic orientation of peptides, resulting in Sammox-driven CRNs that has both anabolic and catabolic reactions at all times. Furthermore, peptides produced from sulfite-fueled Sammox-driven CRNs could catalyze both sulfite-fueled Sammox and Anammox (nitrite reduction coupled with anaerobic ammonium oxidation) reactions. We propose that Sammox-driven CRNs were critical in the creation of life and that Anammox microorganisms that have both Sammox functions are direct descendants of Sammox-driven CRNs.

## INTRODUCTION

The chemistry of life can be thought of as autocatalytic organized chemical reaction networks (CRNs), which involve coupling transformation of six key elements—carbon (C), hydrogen (H), oxygen (O), nitrogen (N), sulfur (S), and phosphorous (P) (Kauffman 1986; Falkowski et al., 2008; Hordijk et al., 2018). We speculate that there might have been a thermodynamic chemical reaction network, which involved C, H, O, N, and S, initiated by exergonic redox reactions, resulting in the development of the proto-metabolic networks (PMNs) in primordial phosphorus-deficient circumstances. The importance of N and S geochemical transformations in the origin of life has been majorly overlooked. A recent study has demonstrated the vital role of sulfur reduction and anaerobic ammonium oxidation in the origin of life (Li et al., 2020a). Phylogenetic distribution and functional grouping of sulfite reductase clusters show that a sulfite reductase, with a coupled siroheme-[Fe_4_-S_4_] cluster, was most likely present in the last universal common ancestor (LUCA) (Crane et al., 1995; Molitor et al., 1998; Dhillon et al., 2005). Nitrite reduction coupled with anaerobic ammonium oxidation (Anammox) might be an ancient metabolism because Anammox organisms could be the first nascent bacterial species (Brochier and Philippe, 2002; van Niftrik and Jetten, 2012), and might be the evolutionary kinds between the three domains of life (Reynaud and Devos, 2011).

Sulfite was richly produced on early Earth from volcanic and hydrothermal sulfur dioxide (Ono et al., 2003; Canfield et al., 2006; Anbar 2008; Falkowski et al., 2008; Moore et al., 2017). Most of the nitrogen species in hydrothermal fluids released from the mantle of the reduced young Earth into the early oceans might have comprised mostly ammonium (Li and Keppler, 2014; Mikhail and Sverjensky, 2014). As a result, spontaneous redox reaction transfers for energy generation and organic molecule synthesis occurred when the prebiotically plausible sulfurous species and ammonium in early Earth hydrothermal environments encountered carbon dioxide. We hypothesize that thermodynamically feasible CRNs containing HCO_3_^−^, H_2_O, NH_4_^+^, SO_3_^2−^/SO_4_^2−^, and HS^-^ (Scheme 1) (Balcerowiak 1985; Amend and Shock, 2001; Fdz-Polanco et al., 2001; Amend et al., 2003; Ma et al., 2009; Schrum et al., 2009; Wang et al., 2013; He et al., 2019; Li et al., 2020b; Wu et al., 2020), might produce PMNs. In this CRNs, sulfur reduction, nitrite reduction coupled with anaerobic ammonium oxidation are all included (Scheme 1; Eq 1, 2, and 3). The reducing power from the reaction (Sammox) could drive the CRNs (Li et al., 2020a).

**Scheme 1.**
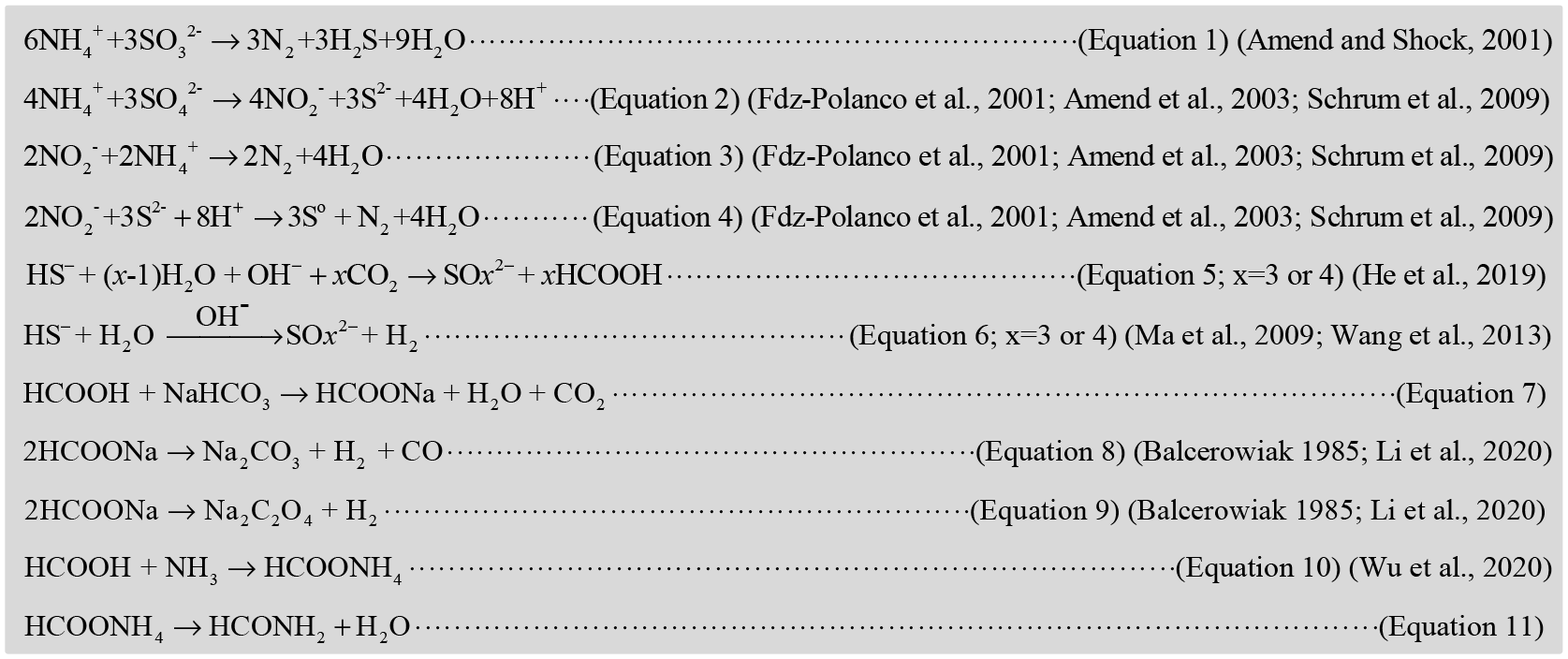
Proposed reactions for the Sammox-driven CRNs. Sammox: equations 1, 2, 3, 4. CO_2_ reduction: equations 6, 7, 8, 9. N activation: equations 10, 11.

The reactions in Sammox-driven CRNs are nonlinear and connected in intricate ways. As a result, we only offer the reaction equation up to the formamide, which acts as a chemical precursor for the production of metabolic and genetic apparatus intermediate (Saladino et al., 2012), (Scheme 1; Eq 11). The Sammox-driven CRNs were nonequilibrium thermodynamic CRNs, which provide energy and material for the development of PMNs. Peptides are the best-known biocatalysts in the cell and the molecular hubs in the origin of life, therefore the PMNs should have at least started with carbon fixation, reductive amination, and continued until peptides production. (Frenkel-Pinter et al., 2020). As a result, prebiotic CRNs, which resulted in the origin of life should have contributed to the autocatalysis of the PMNs. In this study, we demonstrate evidence of peptides production, and the origin of autocatalysis in Sammox-driven CRNs comprising HCO_3_^−^, NH_4_^+^, SO_3_^2−^/SO_4_^2−^ under mild hydrothermal circumstances providing a novel option of the earliest origin of life.

## RESULTS AND DISCUSSION

### Sammox drives peptides formation and its possible pathway

Determination of the likely end product of Sammox-driven CRNs was the primary goal of this study. We detected formate as initial product of Sammox-driven CRNs, and peptides as the end products of sulfite/sulfate-fueled CRNs (Fig. 1 a, b). H_2_S is an essential reductant in abiotic CO_2_ reduction to organics (Wang et al., 2013). The first observation of CO_2_ reduction to formate with H_2_S in a simulated hydrothermal vent system was reported by He et al. (He et al., 2019) (Scheme 1; Eq 5, 6; x = 3 or 4). During this reaction, over 80% S^2−^ was oxidized to SO_3_^2−^ and others to SO_4_^2−^ (Eq 5). The sulfur redox cycle can be obtained through SO_3_^2−^/SO_4_^2−^ reduction to S^2−^ by organic carbon (Wang et al., 2013; He et al., 2019). In this study, the sulfur redox cycle can be obtained through SO_3_^2−^/SO_4_^2−^ reduction to S^2−^ by NH_4_^+^, the Sammox process (Scheme 2). In a different manner, NH_4_^+^ could preserve the organic compounds in Sammox-driven CRNs. Water-gas shift reaction occurred in Sammox-driven CRNs, resulting in the production of active H_2_ and CO (Eq 7, 8, 9) (Balcerowiak 1985; Li et al., 2020b). Active H_2_ was employed to reduce CO_2_, and α-keto acids production (Scheme 2). Formate may react with ammonium to produce ammonium formate (Scheme 1; Eq 10) (Wu et al., 2020). Ammonium formate may be both hydrogen and nitrogen sources for the reductive amination of α-keto acids (Wu et al., 2020). Ammonium formate reacted with pyruvate to produce alanine, glycine, valine, leucine, and serine (Table 1; extended data Fig. 1). Ammonium formate reacted with oxaloacetate to produce aspartate, threonine, and a high level of alanine up to 268 μM (Table 1; extended data Fig. 2). Ammonium formate reacted with α-ketoglutarate to produce proline, arginine, and a high level of glutamate up to 46 μM (Table 1; extended data Fig. 3). Ammonium formate reacted with pyruvate,
oxaloacetate, and α-ketoglutarate to produce alanine, glycine, valine, leucine, serine, glutamate, and aspartate. That is because heating ammonium formate turns it into formamide (Scheme 1, Eq 11), which acts as a chemical precursor for pyruvate, oxaloacetate, and α-ketoglutarate production (Saladino et al., 2012). Succinate, malate, and fumarate can be generated from oxaloacetate, by a reductive dehydroxylation step to yield succinate, and a two-electron–two-proton reduction step to yield malate, a β-elimination of H_2_O to yield fumarate, whereas α-ketoglutarate can be produced by adding a one-carbon unit to succinate (Saladino et al., 2012). Other possible pathways, such as carboxylic acids and amino acids, which could be the products of sodium cyanide and ammonium chloride at 38°C cannot be ruled out (Ruiz-Bermejo et al., 2012). Because heating of formamide at higher temperatures and under acidic circumstances will produce hydrogen cyanide, and these conditions can be met in our experiment (Eq. 12, 13).

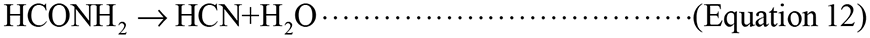

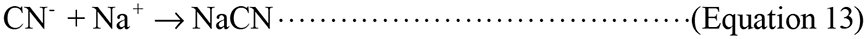

**Scheme 2.**
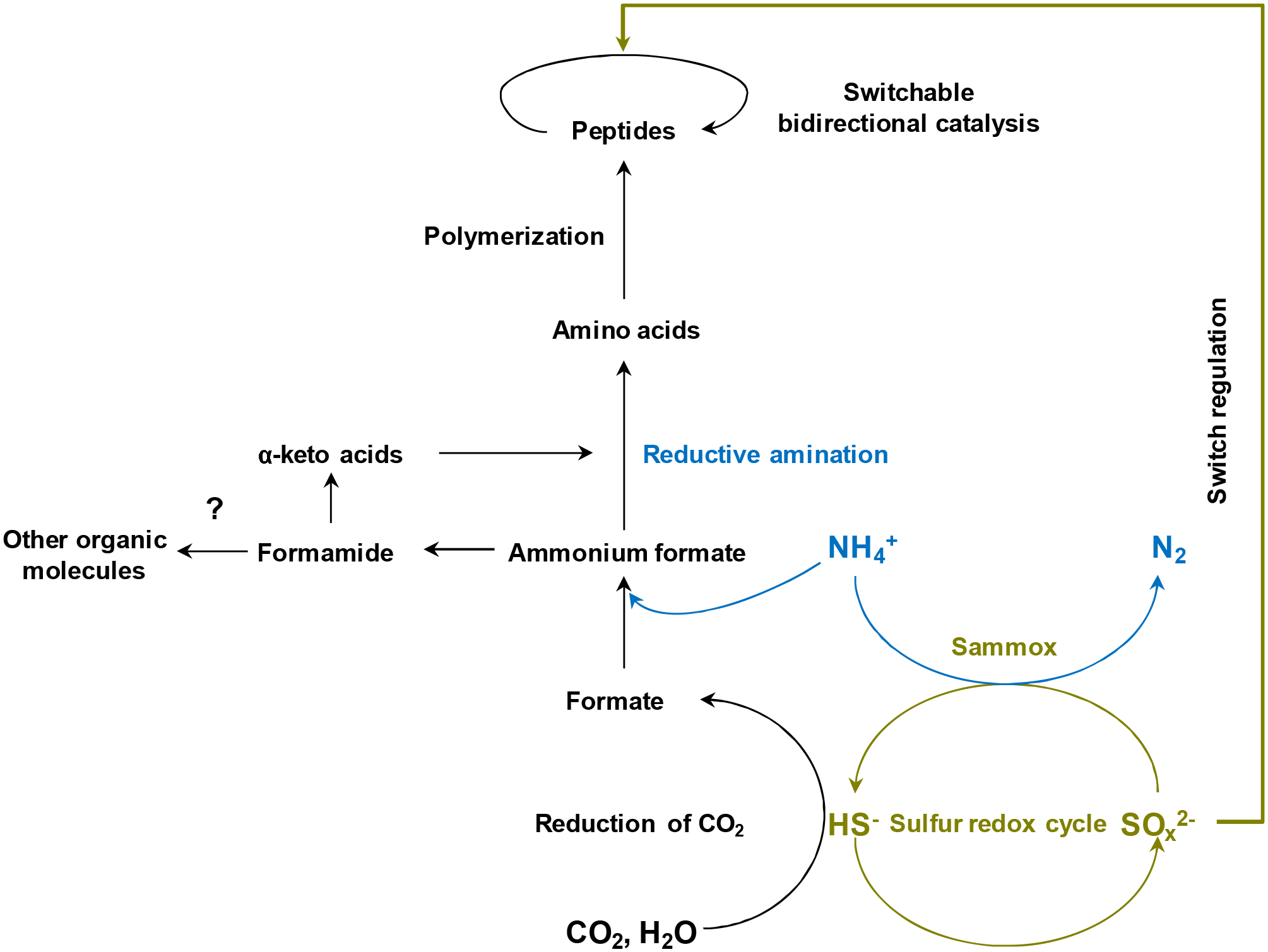
Demonstrates the conceptual model of Sammox-driven coupled transformation of carbon, hydrogen, oxygen, nitrogen, sulfur simultaneously in nonequilibrium thermodynamic environments, initiating the emergence of prebiotic autocatalytic CRNs. Switchable bidirectional catalysis of peptides was regulated by the ratio of sulfite to sulfate in Sammox-driven prebiotic CRNs. x = 3, 4.

**Figure 1.**
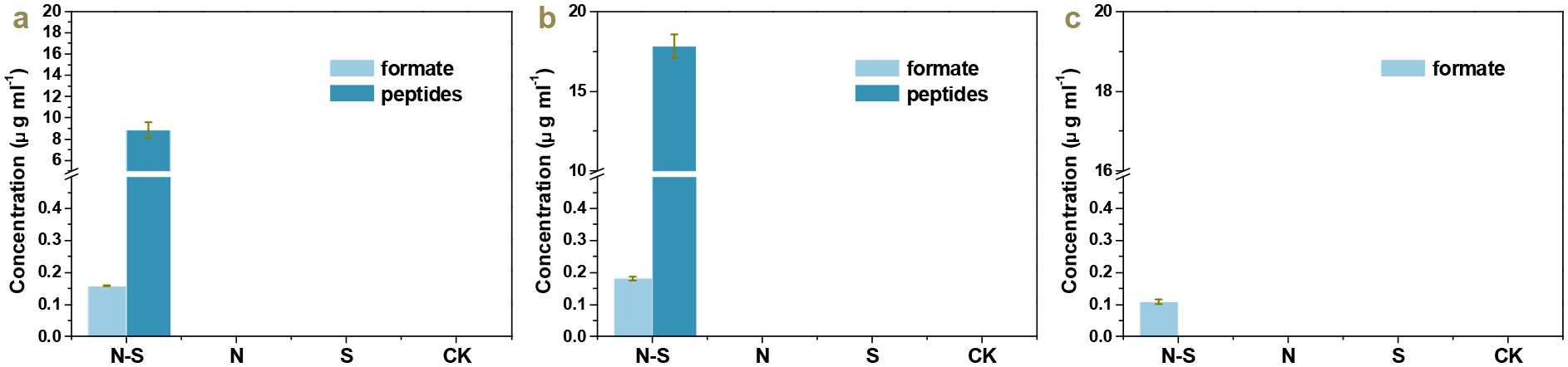
Organic compounds developed from Sammox-driven prebiotic CRNs with bicarbonate as the sole carbon source under mild hydrothermal conditions. Treatments were as follows (a, sulfite-fueled CRNs; b, sulfate-fueled CRNs; c, sulfide-fueled CRNs) from left to right, N-S: ammonium + sulfurous species, N: ammonium, S: sulfurous species, CK. The bar chart indicates the output of formate and peptides in each treatment group. Error bars represent standard deviations of three replicates.

**Figure 2.**
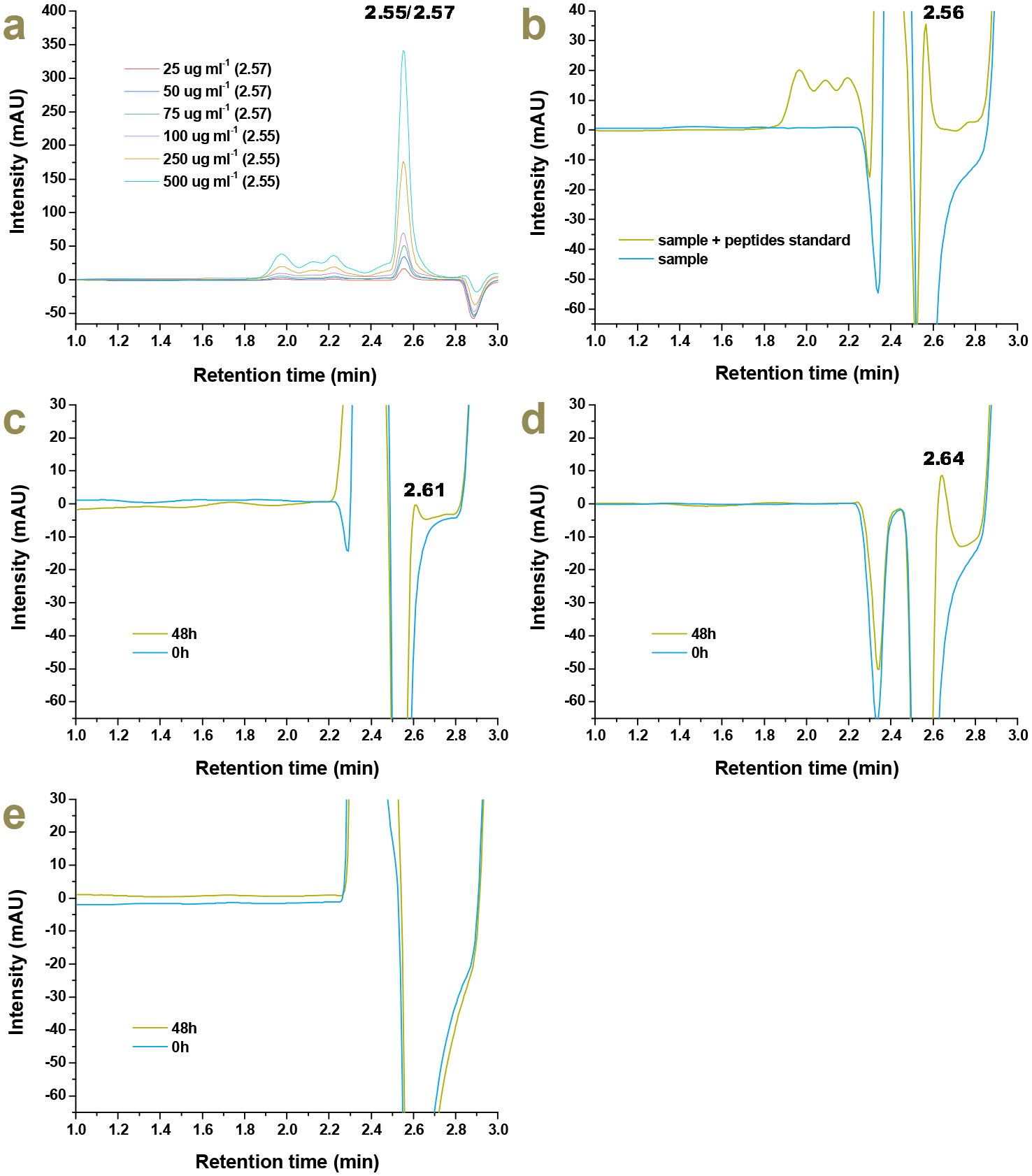
Chromatogram of peptides developed from Sammox-driven CRNs, with bicarbonate as the sole carbon source, under mild hydrothermal circumstances. A, peptides standards (the retention time for each concentration is demonstrated accordingly in brackets); b, a 0.4 ml of 0 h sample of sulfite-fueled Sammox-driven CRNs added into 0.3 ml peptides standards (500 μg ml^−1^); c, sulfite-fueled Sammox-driven prebiotic CRNs; d, sulfate-fueled Sammox-driven prebiotic CRNs; e, sulfide-fueled Sammox-driven prebiotic CRNs. The number illustrates the retention time of peptides peaks.

**Figure 3.**
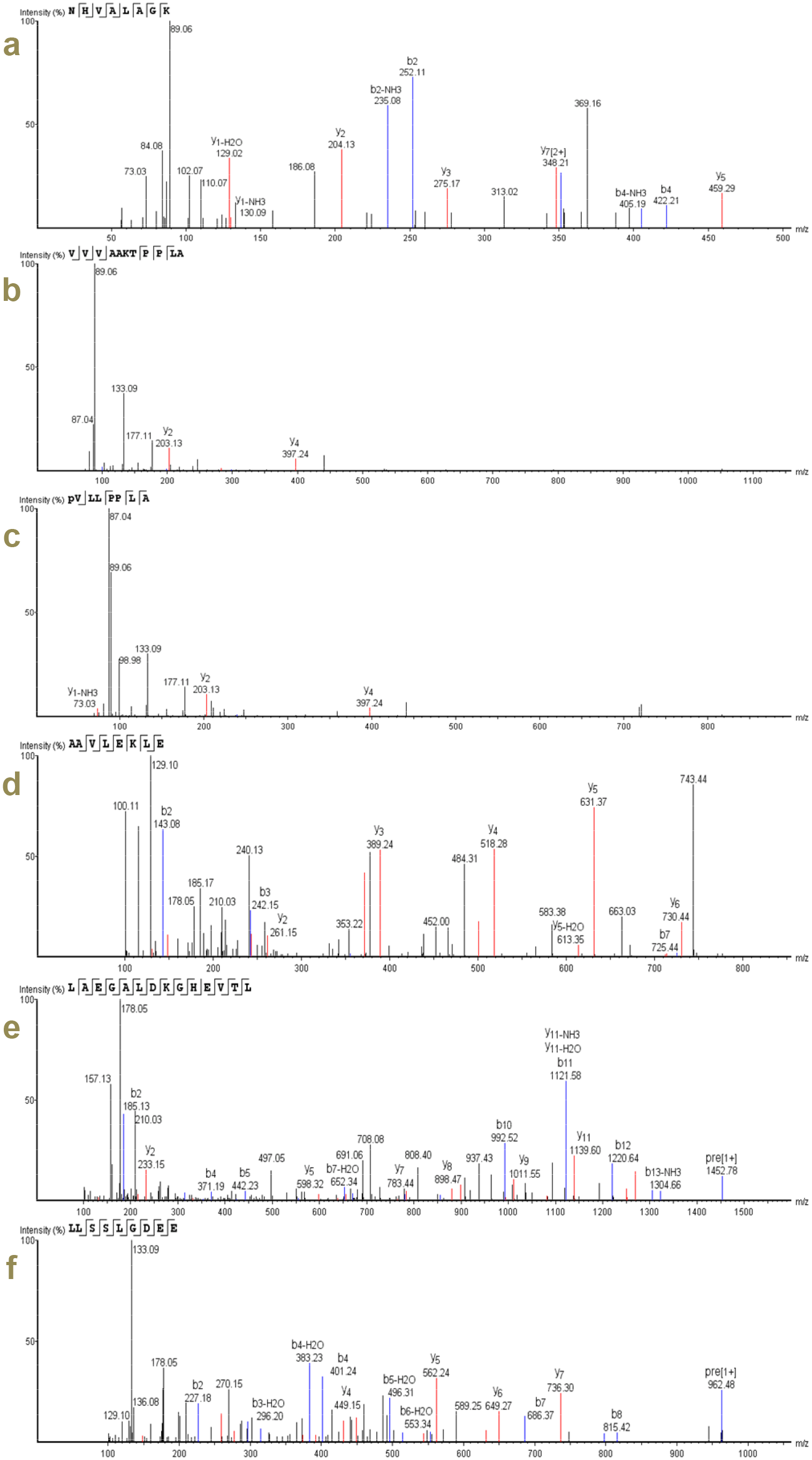
MS/MS spectra of Sammox-driven CRNs-generated peptides. The spectra of selected three peptides a–c were from sulfite-fueled Sammox-driven CRNs. The spectra of selected three peptides d–f were from sulfate-fueled Sammox-driven CRNs.

**Table 1.** Proteinogenic amino acids abundance in peptides, and in reactions of ammonium formate (AF) with α-keto acids. The associated errors are standard deviations of three replicates. ND, not detected.

| Amino acids | Concentrations (μM) | | | | |
| --- | --- | --- | --- | --- | --- |
| | Sulfite-fueled CRNs | Sulfate-fueled CRNs | AF + pyruvate | AF + oxaloacetate | AF + α-ketoglutarate |
| L-Serine | $4.96 \pm 1.35$ | $3.54 \pm 0.65$ | $3.20 \pm 1.17$ | $1.92 \pm 0.90$ | $3.26 \pm 1.67$ |
| Glycine | $3.25 \pm 0.38$ | $2.92 \pm 0.73$ | $3.08 \pm 1.02$ | $2.76 \pm 1.42$ | $5.25 \pm 2.32$ |
| L-Aspartate | $2.42 \pm 0.39$ | $0.97 \pm 0.46$ | $1.03 \pm 0.22$ | $0.77 \pm 0.33$ | $1.79 \pm 0.63$ |
| L-Glutamate | $1.81 \pm 0.26$ | $10.14 \pm 4.17$ | $1.48 \pm 0.24$ | $3.73 \pm 1.12$ | $46.93 \pm 10.92$ |
| L-Alanine | $2.66 \pm 0.38$ | $1.81 \pm 0.52$ | $1.83 \pm 0.18$ | $268.25 \pm 35.69$ | $2.02 \pm 0.21$ |
| L-Asparagine | $0.36 \pm 0.22$ | ND | ND | ND | ND |
| L-Threonine | $0.77 \pm 0.28$ | $0.73 \pm 0.24$ | ND | $1.12 \pm 0.89$ | $0.99 \pm 0.29$ |
| L-Arginine | $0.40 \pm 0.15$ | $0.55 \pm 0.03$ | ND | ND | $0.79 \pm 0.44$ |
| L-Histidine | $0.53 \pm 0.14$ | $0.64 \pm 0.14$ | ND | ND | ND |
| L-Proline | $0.57 \pm 0.05$ | $0.59 \pm 0.09$ | ND | ND | $0.68 \pm 0.24$ |
| L-Lysine | $0.44 \pm 0.05$ | $0.50 \pm 0.21$ | ND | ND | $0.88 \pm 0.31$ |
| L-Tyrosine | $0.53 \pm 0.34$ | ND | ND | ND | ND |
| L-Valine | $0.49 \pm 0.07$ | $0.53 \pm 0.14$ | $0.72 \pm 0.19$ | ND | $0.82 \pm 0.32$ |
| L-Leucine | $0.67 \pm 0.09$ | $0.74 \pm 0.18$ | $0.97 \pm 0.19$ | $0.78 \pm 0.28$ | $1.20 \pm 0.55$ |

However, there is no peptides production in sulfide-fueled CRNs (Fig. 1c), which may require a higher reaction temperature. As sulfate would have been severely limited in the primordial environment, sulfite-fueled CRNs may be viable Sammox-driven CRNs that contribute to the production of peptides under mild primordial conditions (Canfield et al., 2006; Crowe et al., 2014; Moore et al., 2017; Colman et al., 2020). It is worth mentioning that sulfite should be accelerant during reductive amination (Wang et al., 2012). Figure 2 indicates a chromatogram of peptides generated from Sammox-driven prebiotic CRNs. Figure 2b indicates the peak of standard peptides that appeared in similar retention times of target peptides in the sample, verifying the feasibility of the standard peptides. Table 2 demonstrates selected identified peptides produced from Sammox-driven prebiotic CRNs, and Figure 3 representative MS/MS spectra of identified peptides produced from Sammox-driven prebiotic CRNs. We have not detected free amino acids in Sammox-driven CRNs (data not shown). This indicates that the Sammox-driven CRNs facilitate amino acids polymerization, given that carbon disulfide (CS_2_) and carbonyl sulfide (COS), components of the interaction of hydrogen sulfide and carbon dioxide under mild circumstances in an anaerobic aqueous environment (Heinen and Lauwers, 1996), could promote peptide bond formation (Leman et al., 2004, 2015; Frenkel-Pinter et al., 2020).

**Table 2.**
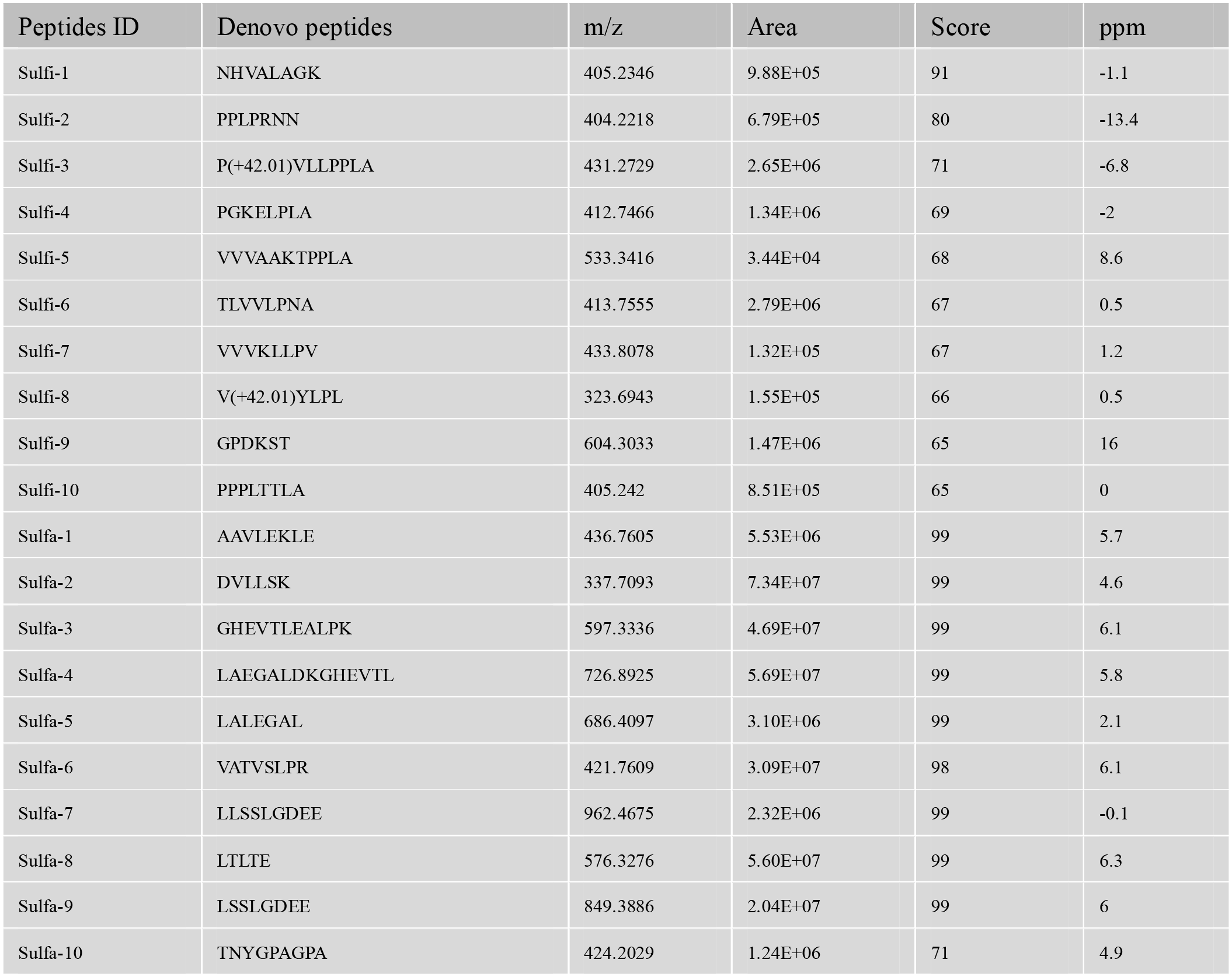
Identified selected peptides in Sammox-driven CRNs. Sulfi, peptides from sulfite-fueled CRNs; Sulfa, peptides from sulfate-fueled CRNs.

We discovered that the peptides comprised 14 proteinogenic amino acids, including L-alanine, glycine, L-valine, L-histidine, L-leucine, L-serine, L-aspartate, L-asparagine, L-lysine, L-glutamate, L-tyrosine, L-threonine, L-proline, and L-arginine (Table 1; extended data Fig. 4, and 5). The 14 proteinogenic amino acids required only a few steps in their metabolism from the incomplete rTCA (Hartman 1975). L-alanine, glycine, L-valine, L-leucine, and L-serine are closely linked to pyruvate. L-aspartate, L-asparagine, L-lysine, L-tyrosine and L-threonine are linked to oxaloacetate. L-glutamate, L-proline, and L-arginine are linked to α-ketoglutarate. Notably, compared to other amino acids, the frequency of the most ancient amino acids—glycine, L-alanine, L-aspartate, and L-glutamate—was relatively high compared to other amino acids (Table 1), which can be explained by existing theories (Wong 2005; Trifonov et al., 2012). The frequency of L-serine was also relatively high, as it is conserved in ancestral ferredoxin (Eck and Dayhoff, 1966). More importantly, the frequency of glycine, L-alanine, L-aspartate, L-glutamate, and L-serine as conserved amino acids is in accordance with the earliest stage of genetic code evolution (Davis 2002).

**Figure 4.**
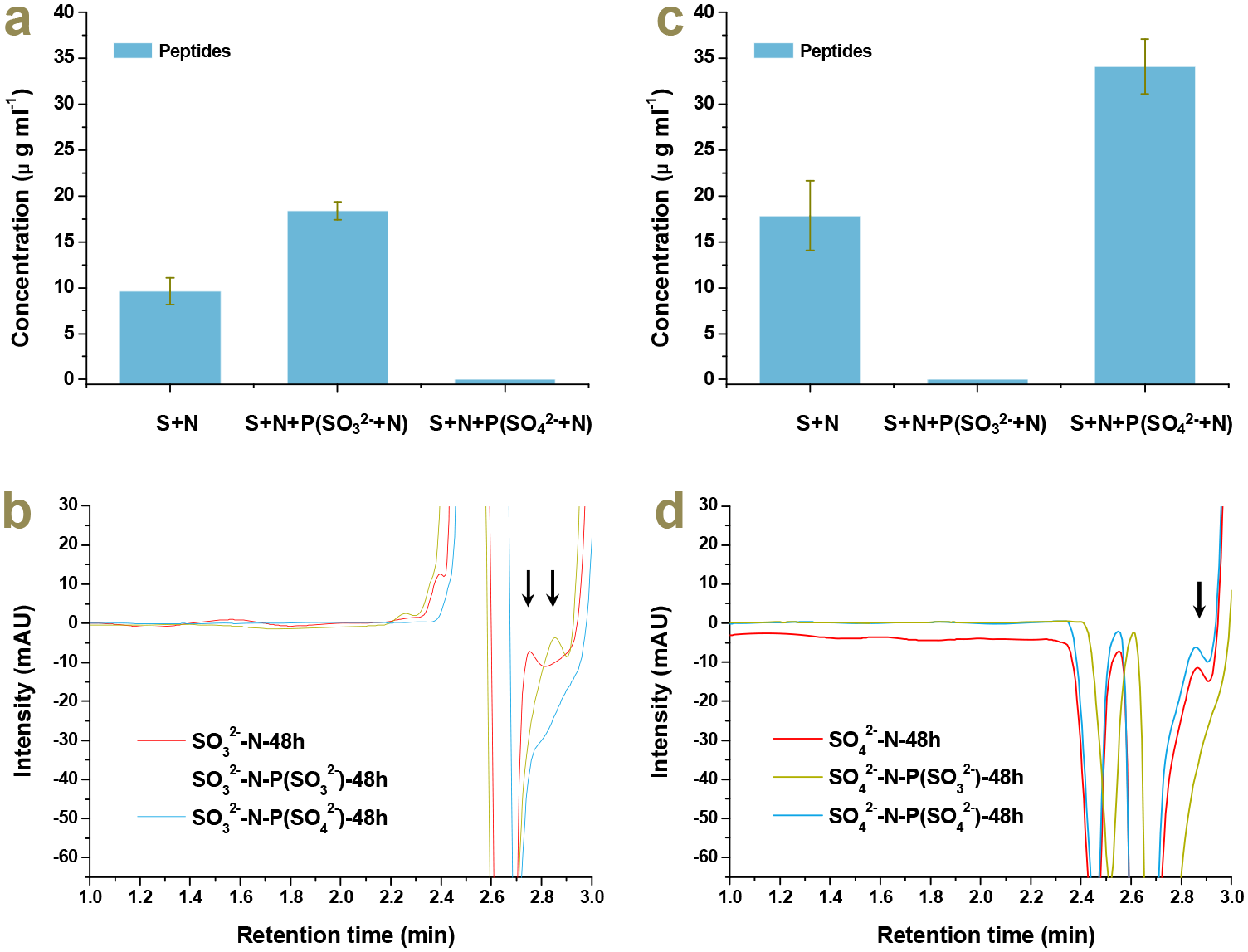
Autocatalysis of the Sammox-driven CRNs under 100°C–70°C after 48 h reaction (a, peptides in sulfite-fueled CRNs; c, peptides in sulfate-fueled CRNs). For a and c, treatments were as follows from left to right: sulfurous species with ammonium; sulfurous species with ammonium plus 1.0 ml products from sulfite-fueled CRNs; sulfurous species with ammonium plus 1.0 ml products from sulfate-fueled CRNs. Chromatogram of peptides generated from Sammox-driven CRNs (b, sulfite-fueled CRNs; d, sulfate-fueled CRNs). For b and d, the red color line indicates the treatment group of sulfurous species with ammonium; olive color line indicates treatment group of sulfurous species with ammonium plus 1.0 ml products from sulfite-fueled CRNs; blue color line indicates treatment group of sulfurous species with ammonium plus 1.0 ml products from sulfate-fueled CRNs. The arrows demonstrate the peaks of peptides. Error bars illustrate standard deviations of three replicates (P < 0.05).

**Figure 5.**
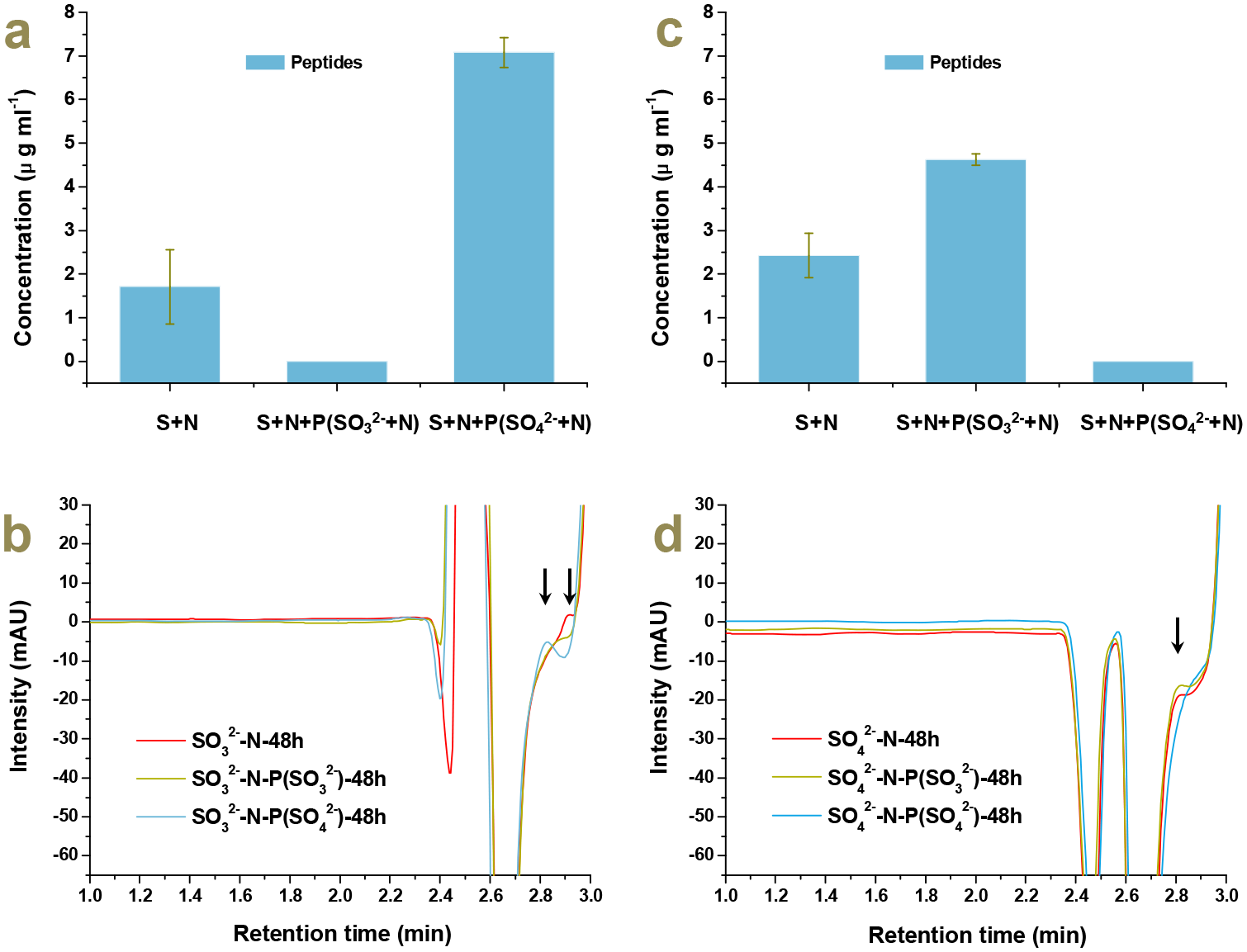
Autocatalysis of the Sammox-driven CRNs under 45°C after 48 h reaction (a, peptides in sulfite-fueled CRNs; c, peptides in sulfate-fueled CRNs). For a and c, treatments were as follows from left to right: sulfurous species with ammonium; sulfurous species with ammonium plus 1.0 ml products from sulfite-fueled CRNs; sulfurous species with ammonium plus 1.0 ml products from sulfate-fueled CRNs. Chromatogram of peptides generated from Sammox-driven CRNs (b, sulfite-fueled CRNs; d, sulfate-fueled CRNs). For b and d, the red color line indicates the treatment group of sulfurous species with ammonium; olive color line indicates treatment group of sulfurous species with ammonium plus 1.0 ml products from sulfite-fueled CRNs; blue color line indicates treatment group of sulfurous species with ammonium plus 1.0 ml products from sulfate-fueled CRNs. The arrows demonstrate the peaks of peptides. Error bars illustrate standard deviations of three replicates (P < 0.05).

### The emergence of autocatalysis in Sammox-driven CRNs

Primordial peptides synthesized by Sammox-driven CRNs should be multifunctional peptides with low substrate specificity; hence, they should be connected mechanistically and evolutionarily. It has been proposed that any sufficiently complex set of polypeptides will inevitably generate reflexively autocatalytic sets of peptides and polypeptides (Kauffman 1986). In 1996, Lee et al. demonstrated that a rationally designed 32-residue α-helical peptide could act autocatalytically in templating its own synthesis by accelerating thioester-promoted amide-bond condensation in neutral aqueous solutions, indicating that the peptide has the possibility of self-replication (Lee et al., 1996). Other studies have proposed that not only do some dipeptides and short peptides have catalytic activities, but even a single proline can have aldolase activity (Sakthivel et al., 2001; Jarvo and Miller, 2002).

In this study, we established series of investigations to confirm whether the Sammox-driven CRNs already autocatalysis, by using Sammox-driven CRNs products to self-catalyze Sammox-driven CRNs. A 1.0 ml Sammox reaction solution (the first round sulfite/sulfate-fueled Sammox) was injected into freshly prepared Sammox reaction solution (the second round sulfite/sulfate-fueled Sammox) as the potential catalyst (Scheme 3 a and b). First, we examined the potential catalysis of products to Sammox-driven CRNs in a variable temperature range, 100°C 24 h–70°C 24 h. Then, we investigated the potential catalysis of products to Sammox-driven CRNs in a selected biologically significant temperature, 45°C 48 h. Surprisingly, products that were generated from sulfite-fueled Sammox reaction could facilitate peptides generation in sulfite-fueled Sammox-driven CRNs at 100°C–70°C, and in sulfate-fueled Sammox-driven CRNs at 45°C; but inhibited peptides generation in sulfate-fueled Sammox-driven CRNs at 100°C–70°C, and in sulfite-fueled Sammox-driven CRNs at 45°C (Scheme 3 a and b; Fig. 4, 5). Products that were generated from sulfate-fueled Sammox reaction, could inhibit peptides generation in sulfite-fueled Sammox-driven CRNs at 100°C–70°C, and in sulfate-fueled Sammox-driven CRNs at 45°C; but facilitate peptides generation in sulfate-fueled Sammox-driven CRNs at 100°C–70°C, and in sulfite-fueled Sammox-driven CRNs at 45°C (Scheme 3 a and b; Fig. 4, 5). As we have demonstrated that there are no free amino acids in Sammox-driven CRNs. The Sammox-driven CRNs-generated peptides are most likely to be the catalysts, which could be complex enough to facilitate the emergence of reflexive autocatalysis, making Sammox-driven CRNs autocatalytic (Scheme 2, 3 c). The peptides might have the reverse catalytic capacity that led to the inhibition phenomenon of peptides production in the corresponding treatment groups. It seemed that peptides presented bidirectional catalysis, forward and reverse catalysis, and always showed the opposite catalytic effect in sulfite- and sulfate-fueled Sammox-driven CRNs, respectively, at selected high and low temperatures, presenting seesaw-like catalytic properties (Scheme 3 d). The catalytic direction of peptides was switchable owing to the ratio of sulfite to sulfate, keeping Sammox-driven CRNs with both anabolic and catabolic reactions at all times. As enzymes are intrinsically bidirectional, a newly evolved enzyme can not be assigned to either autotrophic or heterotrophic metabolism. Enzymes that were able to catalyze the synthesis of CO_2_ to organic macromolecules were principally able to catalyze the degradation of the respective products as well (Gutekunst 2018). So far, we do not know the exact intrinsic mechanisms of seesaw-like catalytic impacts, but it is critical to maintaining the materials balance of Sammox-driven CRNs and the sustainability of Sammox-driven CRNs. This result suggests a common origin of evolutionary metabolism and catabolism since sulfite and sulfate were consistently co-occurring in the Sammox-driven CRNs through the sulfur oxidation cycle (Scheme 2).

**Scheme 3.**
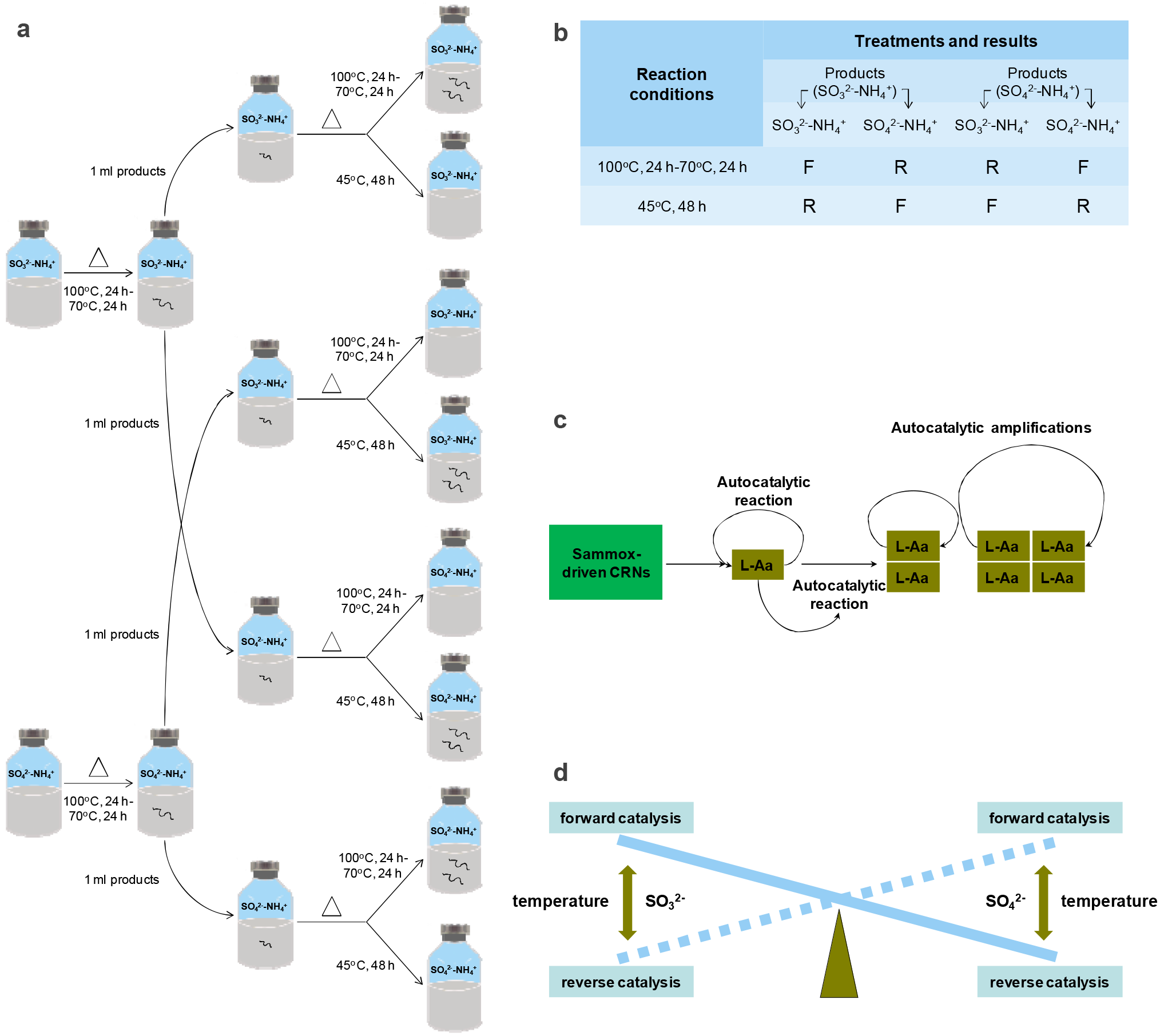
Schematic diagram of detection of autocatalysis in the Sammox-driven CRNs. A and B, experimental procedure and results summary (F: forward catalysis, R: reverse catalysis, : peptides). C, Seesaw-like catalytic impacts of peptides in sulfite and sulfate-fueled Sammox-driven CRNs. D, expected autocatalytic amplification of L-peptides in the Sammox-driven CRNs.

### Relationship between Sammox-driven CRNs and proto-metabolisms

The Sammox-driven CRNs might be PMNs of both autotrophy and heterotrophy. As a result, there may be multiple types of metabolisms stemming from the Sammox-driven CRNs. For example, Sammox, Anammox, formate/hydrogen oxidation coupled with dissimilatory sulfite reduction, dissimilatory nitrite reduction to ammonium (nitrite ammonification), or peptides/protein degradation. If Sammox-driven CRNs could have driven the emergence of LUCA, the direct descendant of LUCA should have similar energy conservation pathways as Sammox-driven CRNs. Accordingly, Planctomycetes could be the first emerging bacterial group, based on analysis of the bacterial phylogeny through rRNA sequences (Brochier and Philippe, 2002). It is also declared that Planctomycetes might be transitional forms between the three domains of life (Reynaud and Devos, 2011), implying a planctobacterial origin of Nomura (eukaryotes, archaea) (Cavalier-Smith and Chao, 2020). The planctomycete *Gemmata obscuriglobus* was the only microorganism capable of protein endocytosis and degradation, implying an intermediate stage between bacteria and eukaryotes (Lonhienne et al., 2010; Acehan et al., 2014). On the other hand, all Anammox microorganisms belong to a monophyletic group, deepest branching inside the phylum Planctomycetes (van Niftrik and Jetten, 2012), and is the only microorganism with membranes comprising both ether-linked lipids (found in archaeal lipids) and ester-linked lipids (found in bacterial and eukaryotic lipids), suggesting a plausible intermediate for the development of the archaeal membrane (Devos and Reynaud, 2010). Therefore, Anammox microorganisms might be the predominant primordial species in Planctomycetes.

A recent study demonstrated the mixotrophic feather of Anammox microorganisms *Candidatus* “*Kuenenia stuttgartiensis”* directly assimilates extracellular formate through the Wood–Ljungdahl pathway instead of oxidizing it completely to CO_2_ (Christopher et al., 2021). Experimental investigation together with genomic evidence has also inferred that Anammox microorganisms can perform reverse metabolism of Anammox, which utilize alternative electron donors to ammonium, such as formate, acetate, and propionate for energy conservation with nitrite or nitrate as electron acceptors (Güven et al., 2005; Strous et al., 2006; Kartal et al., 2007; Christopher et al., 2021). The proto-metabolisms that may allow LUCA to absorb electrons from carbohydrate oxidation and provide a reductant for CO_2_ fixation could have evolved in two connected organisms or a single cell (Gutekunst 2018). A pure culture of Anammox microorganisms is yet to be discovered (Kuenen 2020), implying that Anammox microorganisms may require symbiont to survive. The unique characteristics of Planctomycetes are consistent with the inferences for proto-metabolism, and Anammox microorganisms may be the direct descendant of Sammox-driven CRNs. The functional microorganisms were majorly Anammox microorganisms or Planctomycetes in sulfate-dependent ammonium oxidation environment (Liu et al., 2021), suggesting that the Sammox-driven CRNs can support primordial Anammox metabolism. Indeed, nitrite reduction coupled with ammonium oxidation occurred in Sammox-driven CRNs (Scheme 1, Eq 3).

In this study, we proposed that the peptides generated from sulfite-fueled Sammox-driven CRNs should catalyze the key reactions, such as sulfite-fueled Sammox (Scheme 1, Eq 1) and Anammox reactions (Scheme 1, Eq 3). Dissimilatory sulfite reductase (DsrAB) is closely related to the assimilatory enzyme present in all domains of life and is an enzyme of primordial origin (Wagner et al., 1998; Grein et al., 2013). Because the functional divergence of assimilatory and dissimilatory sulfite reductases preceded the separation of the bacterial and archaeal domains (Crane et al., 1995; Molitor et al., 1998; Dhillon et al., 2005), LUCA most likely had a primordial sulfite reductase. Sulfite respiration required sulfite, formate, or hydrogen all of which were present in the Sammox-driven CRNs (Scheme 1, Figure 1), implying the feasible emergence of sulfite reductase from Sammox-driven CRNs. Sulfite reductases from some sources can catalyze the reduction of both sulfite and nitrite (Crane and Getzoff, 1996), acting as nitrite reductase. As a result, the peptides generated from sulfite-fueled Sammox-driven CRNs may also catalyze Anammox reaction, as nitrite reductase (Nir) is a key enzyme in Anammox reaction (Strous et al., 2006).

Our results demonstrated that peptides that were generated from sulfite-fueled Sammox reaction solutions, could significantly facilitate the consumption of sulfite and ammonium in sulfite-fueled Sammox-driven CRNs (P < 0.05) (Fig. 6 a, b). More sulfate was generated through the sulfur redox cycle (Scheme 2) in the peptides amendment group (Fig. 6 a). We also detected a trace of thiosulfate, as it was reported that thiosulfate can be generated through the sulfur redox cycle (He et al., 2019). Peptides that were generated from sulfite-fueled Sammox reaction solutions could significantly facilitate the consumption of nitrite and ammonium in the Anammox reaction (P < 0.05) (Fig. 6 c). This finding infers an intrinsic relationship between Sammox-driven CRNs and proto-metabolisms, and the common evolutionary origin of Sammox and Anammox. Anammox microorganisms may be the direct descendant of Sammox-driven CRNs. It will be critical to investigate the Sammox function of Anammox microorganisms.

**Figure 6.**
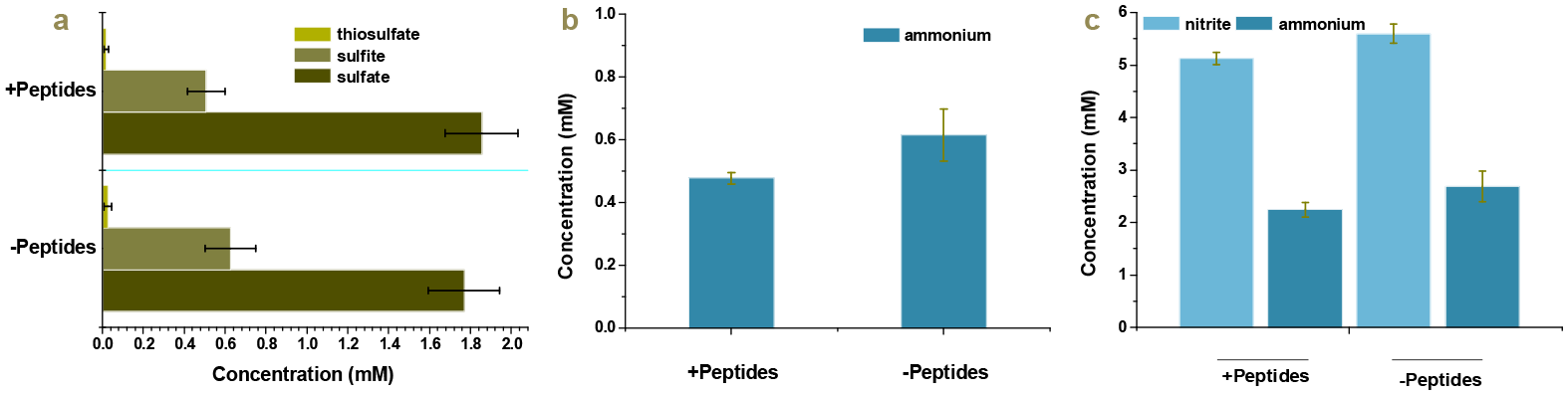
Peptides that were generated from sulfite-fueled Sammox reaction solutions, facilitated the consumption of sulfite and ammonium in sulfite-fueled Sammox-driven CRNs, and nitrite and ammonium in Anammox reaction. Data are all obtained after 48 h of reaction. Error bars indicate standard deviations of three replicates (P < 0.05).

## SIGNIFICANCE

This study reports that the simplest substances—CO_2_, sulfite/sulfate, and ammonium—were converted to peptides in one geological setting by Sammox-driven CRNs which consisted of CO_2_ fixation and reductive amination. Peptides, with 14 proteinogenic amino acids, provide the Sammox-driven CRNs with autocatalysis. Peptides exhibit bidirectional catalysis, with the opposite catalytic effect in sulfite and sulfate-fueled Sammox-driven CRNs, respectively, at both a variable temperature range and a fixed temperature, resulting in seesaw-like catalytic characteristics. The seesaw-like catalytic characteristics of peptides enable Sammox-driven CRNs to maintain both anabolic and catabolic reactions at all times. This result suggests a common origin of primordial metabolism and catabolism since sulfite and sulfate were co-occurring consistently in the Sammox-driven CRNs through the sulfur redox cycle. In addition, peptides generated from sulfite-fueled Sammox-driven CRNs can catalyze both sulfite-fueled Sammox and Anammox reactions, combining the unique characteristics of Anammox microorganisms with the inference of proto-metabolism, Anammox microorganisms with both Sammox functions may be the direct descendant of Sammox-driven CRNs. We infer that Sammox-driven CRNs, under mild conditions, are critical for driving the origin of life.

## MATERIAL AND METHODS

### Chemicals and reagents

All chemical reagents and organic solvents were of analytical grade. Detailed information is as follows: ammonium chloride (>99.5%, CAS number: 12125–02–9), ammonium formate (>97%, CAS number: 540-69-2) were acquired from Sigma–Aldrich, USA. Sodium sulfate (>99.9%, CAS number: 7757-82-6), sodium sulfide nonahydrate (>98%, CAS number: 1313–84–4), acetonitrile (>99.9%, CAS number: 75-05-8), methanesulfonic acid (>99.5%, CAS number: 75–75–2) were purchased from Aladdin, USA. Sodium sulfite (>98.5%, CAS number: 7757–83–7), oxalacetric acid (>98%, CAS number: 328–42–7), 2-Ketoglutaric acid, disodium salt, dehydrate (>99%, CAS number: 305–72–6) were purchased from Acros Organics, Belgium. Pyruvic acid sodium (>98%, CAS number: 113–24–6) was purchased from Amresco, USA. Methanol (>99.9%, CAS number: 67–56–1) was obtained from Tedia, USA. Sodium acetate anhydrate (>99%, CAS number: 127-09–3) was purchased from Sangon Biotech, China. Sodium bicarbonate (>99%, CAS number: 144–55–8) was purchased from Macklin, China. Peptide digest assay standard (Pierce^TM^ Quantitative Colorimetric Peptide Assay, Thermo Scientific, USA) was as standard for quantitative analysis of peptides. All reagents were utilized without further purification unless otherwise noted. Ultrapure water was prepared employing the Millipore purification system (Billerica, MA, USA).

### General procedure for Sammox reactions

A total of 100 mL ultrapure water was introduced into 120 mL serum bottles and sealed with butyl rubber stoppers and aluminum crimp caps. The solution in the serum bottles was autoclaved and cooled at 25°C after being flushed with helium (He) gas (purity = 99.999%). Additional sulfite, sulfate, ammonium, and bicarbonate were introduced into the serum bottles as the “Sammox reaction system.” The above-mentioned ingredients were aseptically introduced to the serum bottles as follows: sodium sulfite, and sulfate (1 mL, 3 mM final concentration), ammonium solution (0.5 mL, 6 mM final concentration for sulfite group; 0.5 mL, 8 mM final concentration for sulfate group), bicarbonate solution (1 mL, 20 mM final concentration). The initial pH value is roughly 8.2. The reaction systems were heated at 100°C in a water bath in the dark for 24 h, maintained at 70°C in the dark for 24 h, and removed from the water bath, and left to cool to room temperature before the investigation was conducted.

This investigation was performed employing the following series of experiments:

(i) 3 mM sulfite/sulfate/sulfide + 6/8 mM NH_4_Cl (for sulfate treatment group) + 20 mM HCO_3_^−^,
(ii) 3 mM sulfite/sulfate/sulfide + 20 mM HCO_3_^−^,
(iii) 6/8 mM NH_4_Cl + 20 mM HCO_3_^−^,
(iv) ultrapure water

### General procedure for the reductive amination of keto acids with HCOONH_4_

To further confirm the feasibility of the reductive amination of keto acids with HCOONH_4_, We employed HCOONH_4_, pyruvic acid, oxaloacetate, and α-ketoglutarate (3.0 mM final concentration), as substrates in hydrothermal reaction systems. Serum bottles were heated at 70°C in a water bath in the dark for 48 h, maintained in the dark, and left to cool to room temperature before investigation. This investigation was conducted employing the following series of experiments:

(i) 3.0 mM HCOONH_4_ + 3.0 mM pyruvic acid.
(ii) 3.0 mM HCOONH_4_ + 3.0 mM oxaloacetate.
(iii) 3.0 mM HCOONH_4_ + 3.0 mM α-ketoglutarate.

### Autocatalysis of the Sammox-driven CRNs

We designed as simple as possible experiments to confirm whether the Sammox-driven CRNs already have autocatalysis. The experimental procedure was conducted in two rounds. In the first round procedure, serum bottles containing Sammox reaction solution were heated at 100°C in a water bath in the dark for 24 h, maintained at 70°C in the dark for another 24 h, and left to cool to room. One milliliter of the first round Sammox reaction solutions (sulfite/sulfate-fueled Sammox) was extracted, and injected into newly prepared Sammox reaction solution (the second round) (sulfite/sulfate-fueled Sammox) as a potential catalyst, respectively. The second round Sammox reaction was executed at two temperature settings 100°C 24h–70°C and 45°C 48 h and allowed to cool to room temperature before sampling. See schemes 3 a, b for a detailed description of the experimental design.

### Peptides facilitate sulfite-fueled Sammox and Anammox reactions

Serum bottles containing sulfite-fueled Sammox reaction solution were heated at 100°C in a water bath in the dark for 24 h, maintained at 70°C in the dark for another 24 h, and left to cool to room. One milliliter of the products was extracted and injected into newly prepared sulfite-fueled Sammox reaction solution (3.0 mM sulfite; 6.0 mM ammonium) and Anammox reaction solution (6.0 mM nitrite; 6.0 mM ammonium), respectively, as a potential catalyst. The reactions were all executed at 100°C, 24h-70°C, 24h, and left to cool to room temperature before sampling for analysis of sulfite, sulfate, thiosulfate, nitrite, and ammonium.

### Sampling analytical methods

#### High-performance liquid chromatography quantitative analysis of peptides

Solution samples were freeze-dried and diluted with ultrapure water to 1.0 mL. The High-performance liquid chromatography system (Agilent LC-1260, USA) comprised a Phenomenex Luna CN 5u column, which is a non-porous analytical column, packed with 5 μm particles (250 mm × 4.6 mm inner diameter, Phenomenex Inc, USA). Mobile phase A comprised 0.05 M sodium acetate, while solvent B was 20% methanol–60% acetonitrile–20% ultrapure water. The samples were investigated utilizing isocratic elution circumstances with an eluent A/B (80:20) for 15 min. The flow rates of the mobile phase and the column temperature were set at 1 mL min^−1^ and 35°C, respectively. The detection wave was UV-214 nm by a diode array detector. Peptides were determined by comparing the retention times against commercially standard peptides. Peptide standard solutions were prepared with concentration gradients of 0, 50, 100, 175, 250, and 400 ug mL^−1^.

#### High-performance liquid chromatography quantitative analysis of thiosulfate

The concentration of thiosulfate was determined employing an Agilent 1260 Infinity HPLC system, equipped with a quaternary pump (Agilent, USA). Thiosulfate was separated with a Zorbax SB-C18 column (150 × 4.6 mm, 5 μm) and detected utilizing a DAD detector at 215 nm. All analyses were performed at 40°C with a flow rate of 1 mL min^−1^. Na_2_HPO_4_ was employed as the solvent. The pH of the solvent was modified with 1.0 M HCl to 8.5. Samples were filtered with 0.45 μm Cosmonice Filters (Millipore, Tokyo, Japan) and immediately injected into the HPLC system (Li et al., 2020).

### Nano LC-MS/MS identification of peptides and amino acids

Solution samples were freeze-dried and diluted with ultrapure water to 1.0 mL. The sample solution was reduced using 10 mM DTT at 56°C for 1 h and alkylated with 20 mM IAA at room temperature, in dark for 1 h. Thereafter, the extracted peptides were lyophilized to almost dryness and resuspended in 2–20 μL of 0.1% formic acid before LC-MS/MS investigation. LC-MS*/*MS investigation was executed on the UltiMate 3000 system (Thermo Fisher Scientific, USA) coupled to a Q Exactive™ Hybrid Quadrupole-Orbitrap™ Mass Spectrometer (Thermo Fisher Scientific, USA). The chromatographic separation of peptides was obtained using a nanocolumn—a 150 μm × 15 cm column—made in-house and packed with the reversed-phase ReproSil-Pur C18-AQ resin (1.9 μm, 100 A, Dr. Maisch GmbH, Germany). A binary mobile phase and gradient were employed at a flow rate of 600 mL min^−1^, directed into the mass spectrometer. Mobile phase A was 0.1% formic acid in the water, and mobile phase B was 0.1% formic acid in acetonitrile. LC linear gradient: from 6%–9% B for 5 min, from 9%–50% B for 45 min, from 50%–95% B for 2 min, and from 95%–95% B for 4 min. The injection volume was 5 μL. MS parameters were set as follows: resolution at 70,000; AGC target at 3e6; maximum IT at 60 ms; the number of scan ranges at 1; scan range at 300 to 1,400 m/z; and spectrum data type was set to profile. MS/MS parameters were set as follows: resolution was set at 17,500; AGC target at 5e4; maximum IT at 80 ms; loop count at 20; MSX count at 1; TopN at 20; isolation window at 3 m/z; isolation offset at 0.0 m/z; scan range at 200 to 2,000 m/z; fixed first mass at 100 m/z; stepped NCE at 27; spectrum data type at profile; intensity threshold at 3.1e4; and dynamic exclusion at 15 s. The raw MS files were investigated and searched against target protein databases, based on the species of the samples utilizing Peaks studio and MaxQuant (1.6.2.10), combined with manual comparison in the UniProt and NCBI databases. The parameters were set as follows: protein modifications were carbamidomethylation (C) (fixed), oxidation (M) (variable), and acetylation (N-term) (variable); enzyme was set to unspecific; the maximum missed cleavages were set to 2; the precursor ion mass tolerance was set to 20 ppm, and MS/MS tolerance was 20 ppm. Only peptides determined with high confidence were chosen for downstream protein determination investigation.

For the analysis of amino acids content, solution samples were freeze-dried, and diluted with ultrapure water to 1.0 ml, followed by acid hydrolysis. A total of 10 μL acid hydrolysate was mixed with 30 μL acetonitrile, vortexed for 1 min, and centrifuged for 5 min at 13,200 r min^−1^ at 4°C. Thereafter, 10 μL of supernatant was introduced to 10 μL water and vortexed for 1 min. Subsequently, 10 μL of the mixture was introduced to 70 μL of borate buffer (from AccQTag kit) and vortexed for 1 min. A total of 20 μL of AccQ Tag reagent (from AccQTag kit) was introduced to the sample, vortexed for 1 min, and the sample was left to stand at ambient temperature for 1 min. Finally, the solution was heated for 10 min at 55°C, and centrifuged for 2 min at 13,200 r min^−1^ and 4°C.

Multiple reaction monitoring investigations were done by utilizing a Xevo TQ-S mass spectrometer. All experiments were conducted in positive electrospray ionization (ESI+) mode. The ion source temperature and capillary voltage were kept constant and set to 150°C and 2 kV, respectively. The cone gas flow rate was 150 L h^−1^ and the desolvation temperature was 600°C. The desolvation gas flow was 1,000 bar. The system was regulated with the analysis software.

### Ion chromatography quantitative analysis of sulfite, sulfate, nitrite, and ammonium

To determine sulfite, sulfate, and nitrite, 1.0 ml of the sample was filtered (0.22 μm) to remove particulates that could interfere with ion chromatography. The ion chromatography system comprised an ICS-5000^+^ SP pump (Thermo Fisher Scientific Inc. Sunnyvale, CA, USA), a column oven ICS-5000^+^ DC, an electrochemical detector DC-5. The ion chromatography column system utilized was a Dionex Ionpac AS11-HC column. The operating condition was with an eluent of 30 mM KOH at a flow rate of 1.0 mL min^−1^. For the determination of ammonium, the ion chromatography column system employed was a Dionex IonpacTM CS 12A column. The operating condition was with an eluent of 20 mM sulphonethane at a flow rate of 1.0 ml min^−1^.

## ACKNOWLEDGMENTS

This research was financially supported by the National Natural Science Foundation of China (General Program Nos. 42077287 and 41571240), and Ningbo Public Welfare project (202002N3101).

## AUTHOR CONTRIBUTIONS

Peng Bao conceived the study, designed and carried out the experiment, and wrote the manuscript. Yu-Qin He and Guo-Xiang Li carried out experiments and analysis. Peng Bao, Yu-Qin He, Guo-Xiang Li, Hui-En Zhang, and Ke-Qing Xiao contributed to interpreting the data. We thanks for Jun-Yi Zhao, Kun Wu, Juan Wang, Xiao-Yu Jia for carrying out sample analysis. Peng Bao wish to dedicate this research to the memory of Mr. Xian-Ming Bao for his kind encouragement and supporting.

