## supplemental figures 1-5 for "A thermodynamic chemical reaction network drove autocatalytic prebiotic peptides formation"

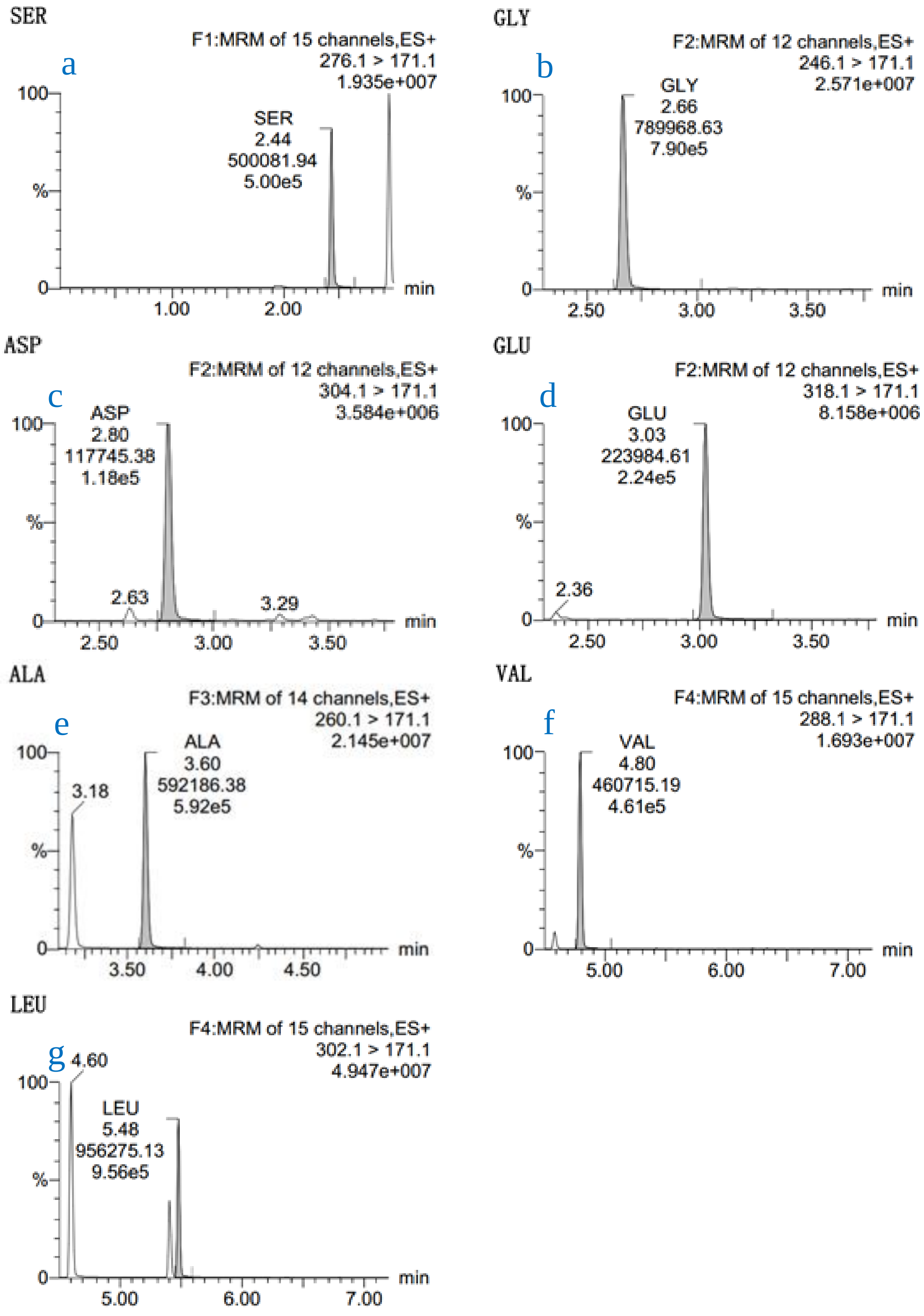


Extended data Figure 1. Chromatogram of proteinogenic amino acids in reactions of ammonium formate (AF) with pyruvate. From a to g: a, Serine; b, Glycine; c, Aspartate; d, Glutamate; e, Alanine; f, Valine; g, Leucine.

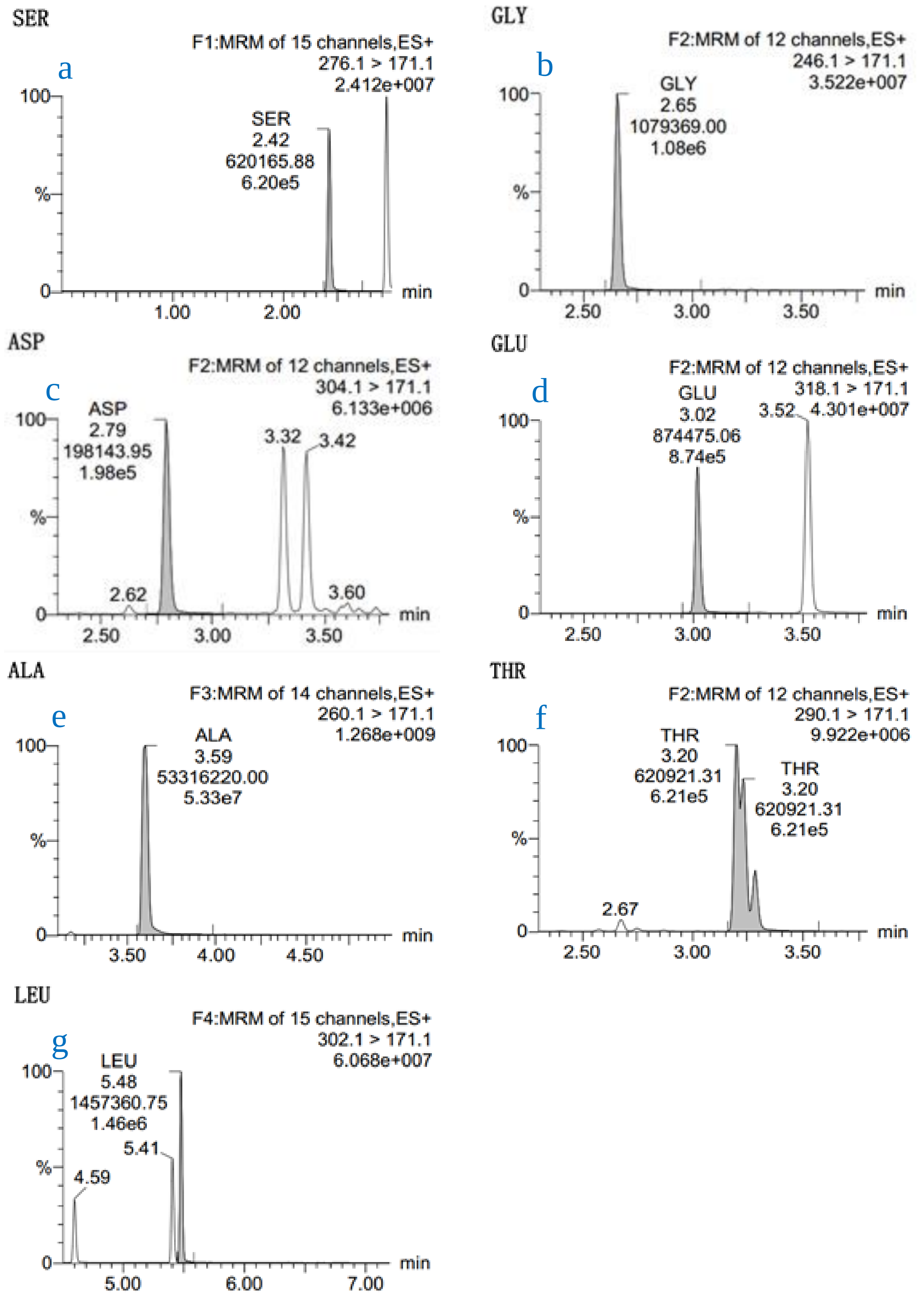


Extended data Figure 2. Chromatogram of proteinogenic amino acids in reactions of ammonium formate (AF) with oxaloacetate. From a to g: a, Serine; b, Glycine; c, Aspartate; d, Glutamate; e, Alanine; f, Threonine; g, Leucine.

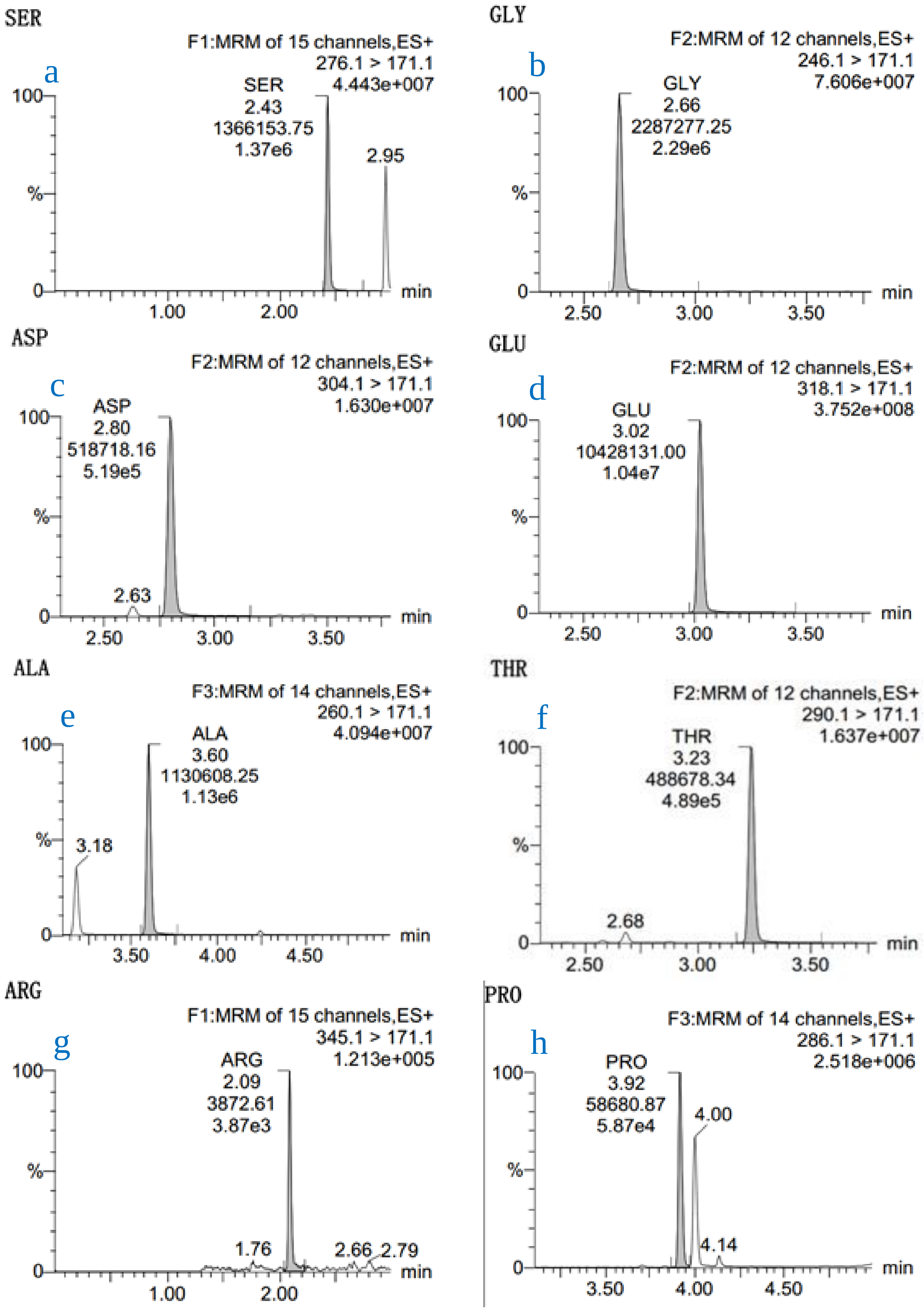
SER
a
F1:MRM of 15 channels,ES+
276.1 > 171.1
4.443e+007
SER
2.43
1366153.75
1.37e6
2.95
100
%
0
1.00
2.00
min
GLY
b
F2:MRM of 12 channels,ES+
246.1 > 171.1
7.606e+007
GLY
2.66
2287277.25
2.29e6
100
%
0
2.50
3.00
3.50
min
ASP
c
F2:MRM of 12 channels,ES+
304.1 > 171.1
1.630e+007
ASP
2.80
518718.16
5.19e5
2.63
100
%
0
2.50
3.00
3.50
min
GLU
d
F2:MRM of 12 channels,ES+
318.1 > 171.1
3.752e+008
GLU
3.02
10428131.00
1.04e7
100
%
0
2.50
3.00
3.50
min
ALA
e
F3:MRM of 14 channels,ES+
260.1 > 171.1
4.094e+007
ALA
3.60
1130608.25
1.13e6
3.18
100
%
0
3.50
4.00
4.50
min
THR
f
F2:MRM of 12 channels,ES+
290.1 > 171.1
1.637e+007
THR
3.23
488678.34
4.89e5
2.68
100
%
0
2.50
3.00
3.50
min
ARG
g
F1:MRM of 15 channels,ES+
345.1 > 171.1
1.213e+005
ARG
2.09
3872.61
3.87e3
1.76
2.66
2.79
100
%
0
1.00
2.00
min
PRO
h
F3:MRM of 14 channels,ES+
286.1 > 171.1
2.518e+006
PRO
3.92
58680.87
5.87e4
4.00
4.14
100
%
0
3.50
4.00
4.50
min

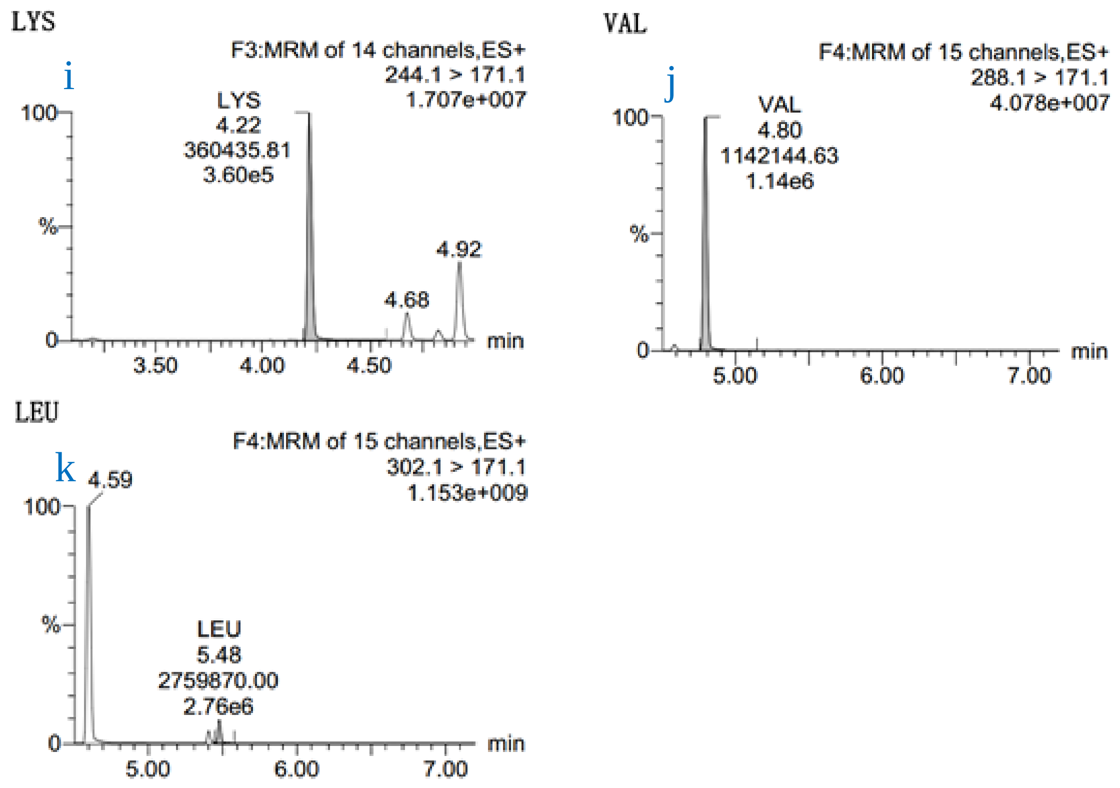


Extended data Figure 3. Chromatogram of proteinogenic amino acids in reactions of ammonium formate (AF) with α-ketoglutarate. From a to k: a, Serine; b, Glycine; c, Aspartate; d, Glutamate; e, Alanine; f, Threonine; g, Arginine; h, Proline; i, Lysine; j, Valine; k, Leucine.

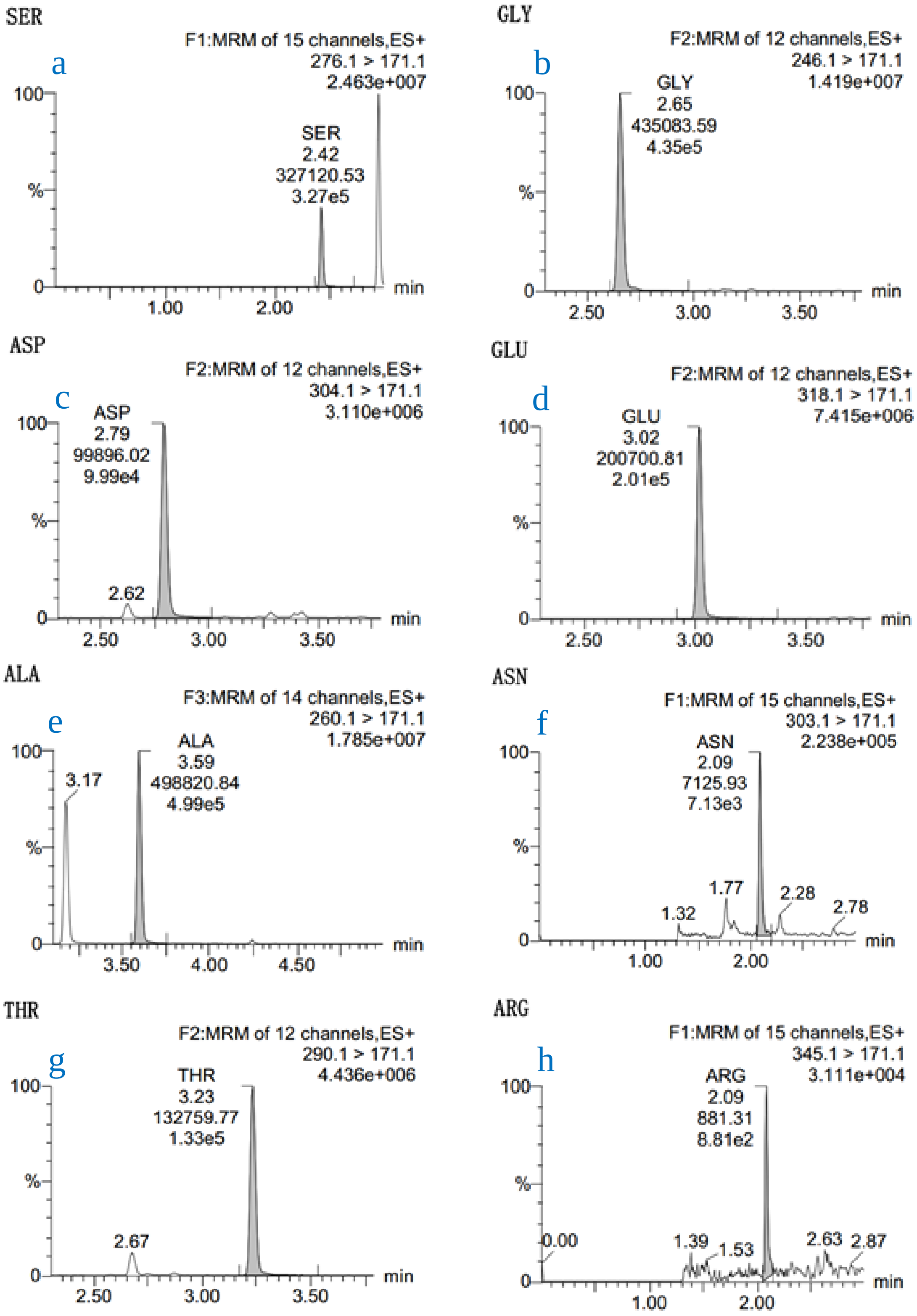
SER
a
F1:MRM of 15 channels,ES+
276.1 > 171.1
2.463e+007
SER
2.42
327120.53
3.27e5
GLY
b
F2:MRM of 12 channels,ES+
246.1 > 171.1
1.419e+007
GLY
2.65
435083.59
4.35e5
ASP
c
F2:MRM of 12 channels,ES+
304.1 > 171.1
3.110e+006
ASP
2.79
99896.02
9.99e4
2.62
GLU
d
F2:MRM of 12 channels,ES+
318.1 > 171.1
7.415e+006
GLU
3.02
200700.81
2.01e5
ALA
e
F3:MRM of 14 channels,ES+
260.1 > 171.1
1.785e+007
3.17
ALA
3.59
498820.84
4.99e5
ASN
f
F1:MRM of 15 channels,ES+
303.1 > 171.1
2.238e+005
ASN
2.09
7125.93
7.13e3
1.32
1.77
2.28
2.78
THR
g
F2:MRM of 12 channels,ES+
290.1 > 171.1
4.436e+006
THR
3.23
132759.77
1.33e5
2.67
ARG
h
F1:MRM of 15 channels,ES+
345.1 > 171.1
3.111e+004
ARG
2.09
881.31
8.81e2
0.00
1.39
1.53
2.63
2.87
100
%
0
min
1.00
2.00
2.50
3.00
3.50
4.00
4.50

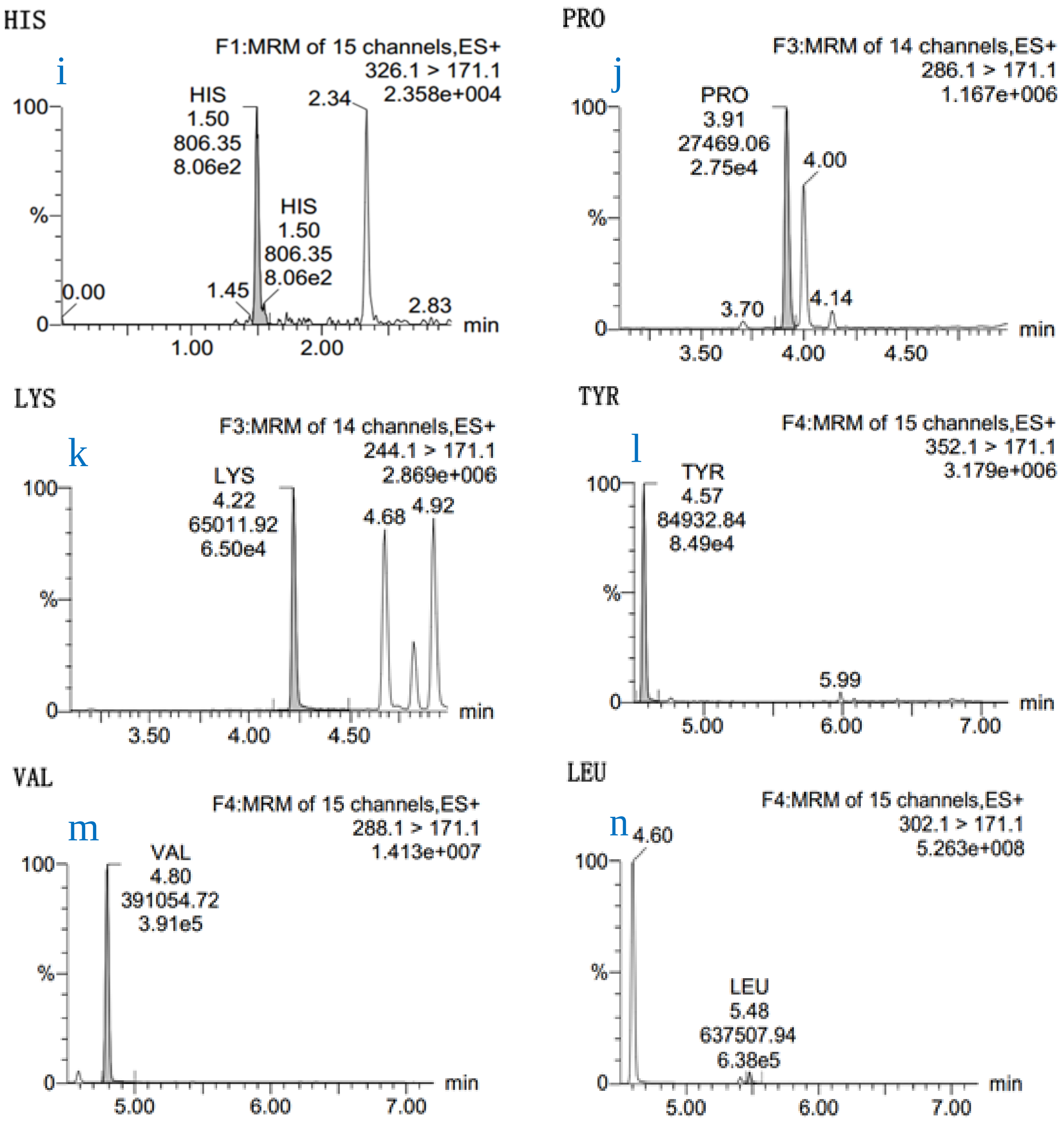


Extended data Figure 4. Chromatogram of proteinogenic amino acids in peptides generated from Sulfite-fueled CRNs. From a to n: a, Serine; b, Glycine; c, Aspartate; d, Glutamate; e, Alanine; f, Asparagine; g, Threonine; h, Arginine; i, Histidine; j, Proline; k, Lysine; l, Tyrosine; m, Valine; n, Leucine.

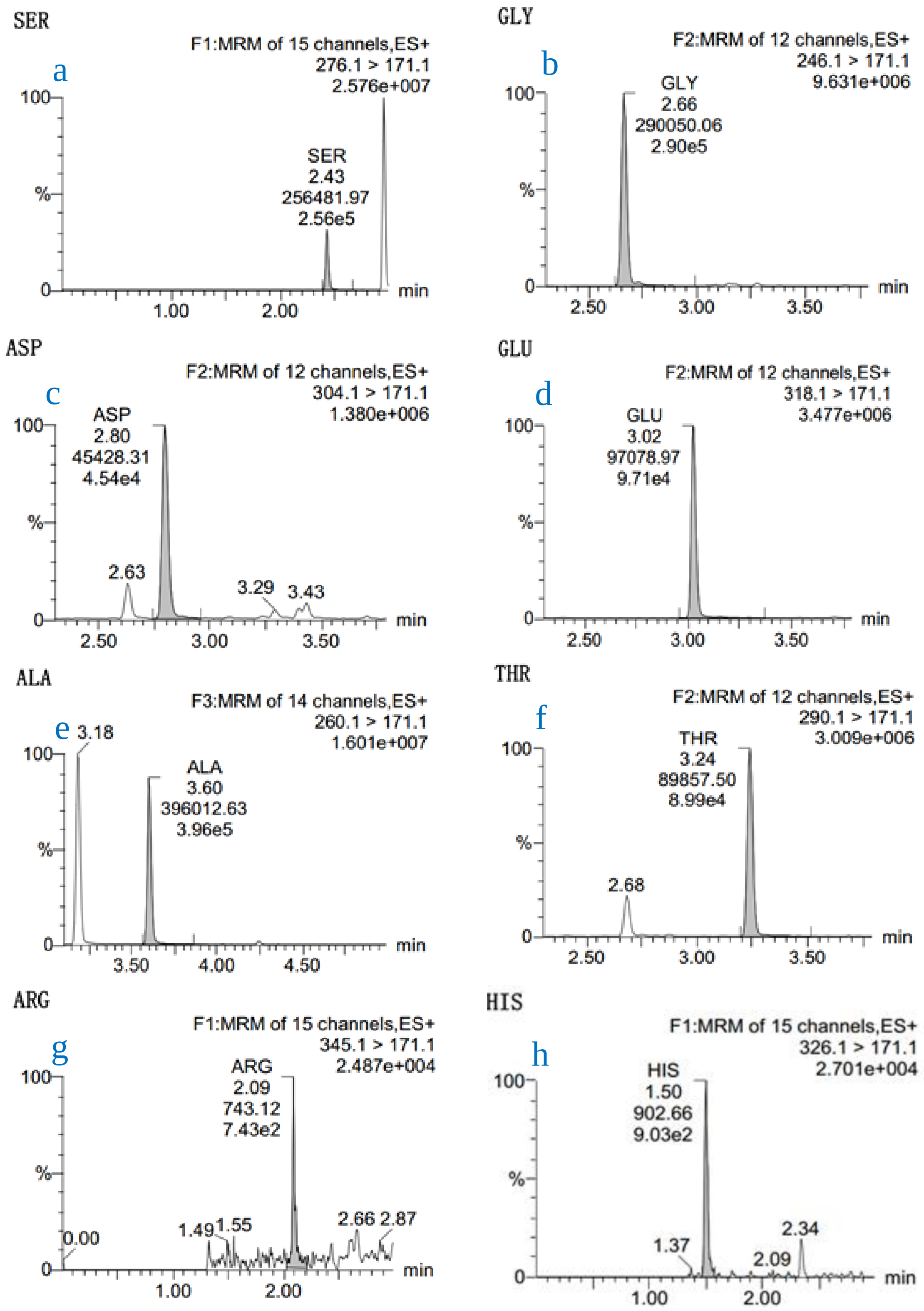
SER
a
F1:MRM of 15 channels,ES+
276.1 > 171.1
2.576e+007
SER
2.43
256481.97
2.56e5
GLY
b
F2:MRM of 12 channels,ES+
246.1 > 171.1
9.631e+006
GLY
2.66
290050.06
2.90e5
ASP
c
F2:MRM of 12 channels,ES+
304.1 > 171.1
1.380e+006
ASP
2.80
45428.31
4.54e4
2.63
3.29
3.43
GLU
d
F2:MRM of 12 channels,ES+
318.1 > 171.1
3.477e+006
GLU
3.02
97078.97
9.71e4
ALA
e
F3:MRM of 14 channels,ES+
260.1 > 171.1
1.601e+007
3.18
ALA
3.60
396012.63
3.96e5
THR
f
F2:MRM of 12 channels,ES+
290.1 > 171.1
3.009e+006
THR
3.24
89857.50
8.99e4
2.68
ARG
g
F1:MRM of 15 channels,ES+
345.1 > 171.1
2.487e+004
ARG
2.09
743.12
7.43e2
0.00
1.49
1.55
2.66
2.87
HIS
h
F1:MRM of 15 channels,ES+
326.1 > 171.1
2.701e+004
HIS
1.50
902.66
9.03e2
1.37
2.09
2.34
100
%
0
min

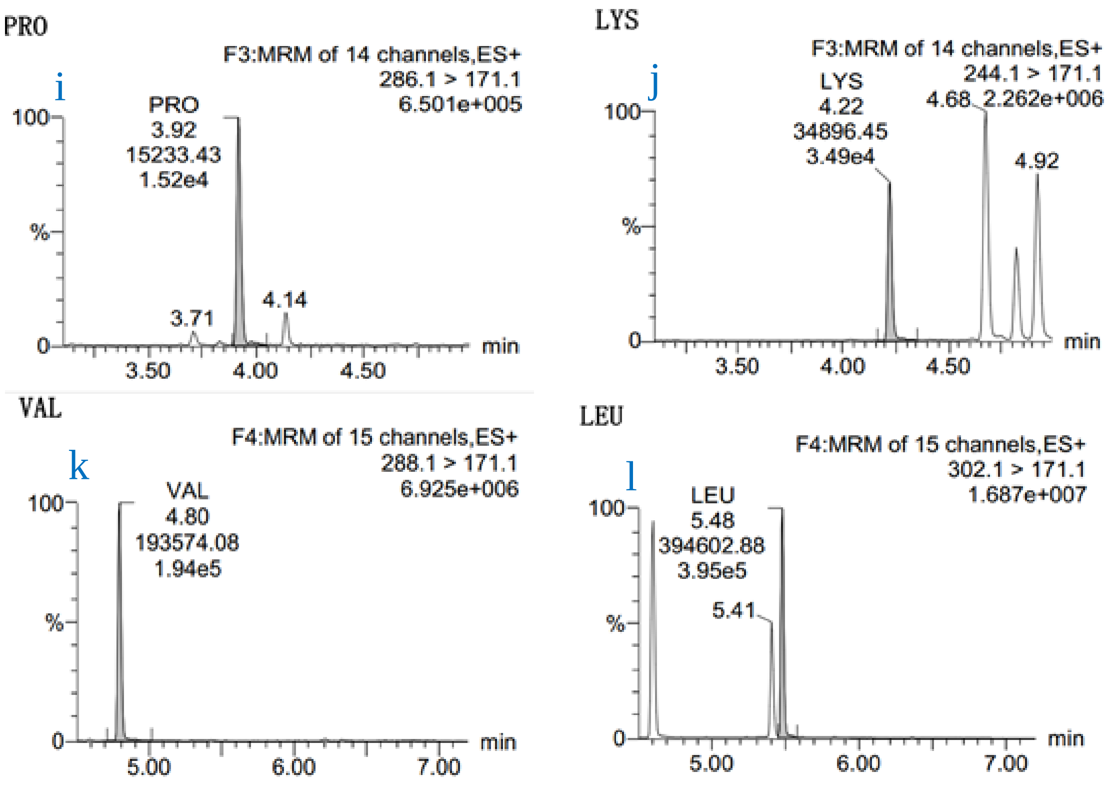


Extended data Figure 5. Chromatogram of proteinogenic amino acids in peptides generated from Sulfate-fueled CRNs. From a to l: a, Serine; b, Glycine; c, Aspartate; d, Glutamate; e, Alanine; f, Threonine; g, Arginine; h, Histidine; i, Proline; j, Lysine; k, Valine; l, Leucine.
